# Building integrated representations through interleaved learning

**DOI:** 10.1101/2021.07.29.454337

**Authors:** Zhenglong Zhou, Dhairyya Singh, Marlie C. Tandoc, Anna C. Schapiro

## Abstract

Inferring relationships that go beyond our direct experience is essential for understanding our environment. This capacity requires either building representations that directly reflect structure across experiences as we encounter them, or computing the indirect relationships across experiences as the need arises. Building structure directly into overlapping representations allows for powerful learning and generalization in neural network models, but building these so-called *distributed representations* requires inputs to be encountered in interleaved order. We test whether interleaving similarly facilitates the formation of representations that directly integrate related experiences in humans, and what advantages such integration may confer for behavior. In a series of behavioral experiments, we present evidence that interleaved learning indeed promotes the formation of representations that directly link across related experiences. As in neural network models, interleaved learning gives rise to fast and automatic recognition of item relatedness, affords efficient generalization, and is especially critical for inference when learning requires statistical integration of noisy information over time. We use the data to adjudicate between several existing computational models of human memory and inference. The results demonstrate the power of interleaved learning and implicate the formation of integrated, distributed representations that support generalization in humans.

## Introduction

Adaptive behavior often requires inferring relationships that go beyond what we have directly experienced. For example, we can infer an efficient route through a city without having traveled the particular path before. There are two broad classes of theories for how we achieve this, which emphasize either discovering and storing these indirect relationships during initial encoding or computing them as needed at the time of a decision (Zeithamova et al., 2012). The retrieval perspective assumes that the representations of indirectly related experiences are kept separate, but that these distinct representations can then interact during retrieval to support inference and generalization (Banino et al., 2016). The advantage of the retrieval-focused representational scheme is that it allows flexibility in how the representations are used; it retains the details of individual elements but allows for these elements to interact and recombine to discover relationships as needed at the time of a decision (Kumaran & McClelland 2012). The disadvantages are that this recombination process takes time and computation at retrieval (Daw & Dayan 2014), and this kind of storage does not scale well (Hinton 1984). The encoding perspective argues that we build representations that directly reflect structure across experiences during initial learning, such that related experiences are built as integrated, overlapping memory representations (Shohamy & Wagner 2008; Eichenbaum et al., 1999; Eichenbaum 2004; Howard et al., 2005; Gluck & Myers 1993). The advantages of this form of “cached” representation are that it scales better and that inference and generalization at retrieval is much more efficient and automatic (Hinton 1986; O’Reilly 1998), but there is a cost in flexibility.

One especially powerful example of the encoding-based representation is the *distributed representation* used in the hidden layers of neural network models. In this form of representation, “each entity is represented by a pattern of activity distributed over many computing elements, and each computing element is involved in representing many different entities” (Hinton, 1984). These representations have proven crucial for modern machine learning models to effectively represent the structure and statistical regularities of complex environments (Hassabis et al., 2017). However, a fundamental limitation of this kind of representation is that it is highly susceptible to interference during distribution shift, that is, in changing environments where exposure to one set of information is mostly completed before beginning the next set (McCloskey & Cohen 1989; McClelland et al., 2020). Consider *blocked* exposure to a set of associations AB, followed by a related set BC. To the extent that the representations of A and C overlap, learning BC will tend to overwrite the connections that had been used for encoding AB (known as “catastrophic interference”). When exposure is *interleaved*, such that AB and BC alternate, their representations can be carefully built up without interference (McClelland et al., 1995); with each input, there is a small amount of interference to other related inputs, but when those related inputs are soon presented again, they regain some lost ground. This delicate back-and-forth allows representations to grow to reflect the full structure of the environment. After this process of interleaved learning, generalization from A to C is then natural and instantaneous, as A and C directly overlap in their representation.

This order-induced interference problem is avoided in retrieval-based models that build distinct representations at encoding, also known as *localist* or *pattern-separated* representations. Keeping representations separate during encoding is very useful for rapid episodic learning without interference, and indeed there is strong evidence for their use towards this purpose in the hippocampus (O’Keefe 1976; Norman & O’Reilly 2003; Kumaran & McClelland 2012; Hainmueller & Bartos 2020). However, as outlined above, this strategy likely limits the efficiency and power of inference and generalization. There is evidence, on the other hand, that relationships are directly encoded in neural patterns throughout the neocortex in the form of distributed representations (Yamins et al., 2014; McClelland et al., 1995; Desimone et al., 1984), but these are believed to form quite slowly, over days, months, and years.

Could distributed representations be supporting our prodigious ability to make inferences and generalize on a faster timescale, over minutes and hours (e.g., finding a good route through a building on your first day encountering it)? There is evidence for the rapid formation of integrated (in addition to separated) representations in the hippocampus (Schlichting & Preston 2015), but it is unclear whether these integrated representations take on the form of the distributed representations found in neural network models, and how they trade off against the well-known pattern-separated representations. Here we use some of the key behavioral signatures of distributed representations — a reliance on interleaved learning and affordance of efficient inference and generalization — to test for their rapid formation. Our predictions are motivated by our neural network model of the hippocampus (Schapiro et al., 2017) which instantiates the hypothesis that a particular circuit, the monosynaptic pathway connecting entorhinal cortex to subfield CA1, rapidly develops distributed representations that allow the hippocampus to detect regularities across novel experiences on the timescale of minutes (Schapiro et al., 2012). The model also simulates the formation of localist representations in subfields CA3 and DG, differentiating individual episodes to avoid interference. We refer to the model as C-HORSE: Complementary Hippocampal Operations for Representing Statistics and Episodes. The model is consistent with neural data: though the hippocampus is known for its pattern-separated representations, rodent work indicates that these reside mainly in the DG and CA3 subfields, while representations appear more overlapping in CA1 (e.g., Leutgeb et al., 2004). Our model thus motivates assessing the formation of distributed representations across the timescale of one human experimental session in the context of a hippocampally-dependent task.

We adopted the hippocampally-dependent associative inference task (Bunsey & Eichenbaum, 1996; Preston et al., 2004), in which participants learn novel object pairs AB and BC and are then tested on the unobserved indirect AC relationship. The basic associative inference task can be solved using either an encoding or retrieval-based strategy (Zeithamova et al., 2012), but here we manipulated whether pairs were learned in interleaved vs. blocked order and tested whether interleaved learning led to behavioral signatures of distributed representations, a particular and powerful flavor of the encoding-based integrated representation. We predicted that, first, interleaved learning should promote implicit, automatic association between A and C at test, whereas blocked learning would require reliance on a localist style representation, requiring additional, and perhaps more explicit, computation to achieve successful inference. Second, integrated, distributed representations should more readily promote generalization: Acquiring new knowledge about A should immediately transfer to C by virtue of their shared neural substrate (Hinton, 1986). Lastly, distributed representations should exhibit more robust graded sensitivity to statistical regularities (Rogers & McClelland 2004), which is not an inherent property of alternative kinds of encoding-based integrated representations (Banino et al., 2016).

There have been many prior studies manipulating order of exposure of information, especially in the category learning literature. While interleaved exposure benefits learning in many variants of category learning tasks (Carvalho & Goldstone, 2015), blocked exposure sometimes appears more beneficial (Flesch et al., 2018; Schlichting et al., 2015; Carvalho & Goldstone, 2015). These findings often appear to hinge on attention being drawn to category similarities and differences across adjacent trials, which is a distinct mechanism from those producing order effects in standard neural network models. We thus endeavored to design a study that would avoid these attentional effects. In our paradigm, back-to-back trials always (in both conditions) contain unrelated pairs such that attention to specific features of back-to-back trials neither benefits nor harms learning, allowing us to isolate effects of order apart from sequential attention.

Each participant in our experiments learned some sets of AB and BC pairs in interleaved order, and other sets in blocked order. If humans learn representations similarly to distributed neural network models, only interleaved exposure should result in behavior that shows the properties outlined above. We provide, to our knowledge, the first evidence that interleaved exposure results in such behaviors. The results indicate a true effect of order, as opposed to a simple influence of time or of trial-by-trial attention modulation. The results are supported by simulations contrasting the behavior of different classes of computational models — those using distributed representations, localist, or both, with the data best explained by the presence of both kinds of representations. Together, the results reveal the benefits of interleaved learning and suggest that humans rapidly form integrated, distributed representations at encoding — likely in addition to more separated representations — in the service of efficient inference.

## Experiments 1a-c

In the first three experiments (Exps 1a-c), each participant learned triads of direct AB and BC associations, with the direct pairs for half of the triads presented in interleaved order and the other half in blocked order (Fig. 1a). After this learning phase, participants completed two tasks that probed their memory for object associations: a speeded recognition task, in which they quickly identified whether a pair of objects shown on the screen was presented during learning (Fig. 1b), followed by an explicit inference task, in which they more slowly and deliberately identified indirect associations (Fig. 1c). In the speeded recognition task, for all pairs of objects except those forming an AB or BC pair, the correct answer was to indicate that they were *not* shown together during learning. This decision could be challenging for the indirectly related AC pairs, as they were associated but not directly studied together. Our main prediction was that this would be especially challenging in the interleaved condition. If interleaving better facilitates overlapping representations, there would be more difficulty rejecting ACs, resulting in increased response time. This confusion could even result in false alarms for interleaved ACs (deciding they had viewed A and C directly together when they had not). For the blocked condition, we predicted a reliance on separated, localist representations of AB and BC, which should prevent this kind of confusion on AC recognition trials. For the subsequent inference task where participants are encouraged to explicitly search for indirect links amongst objects and have enough time to do so, we predicted no difference between AC inference in the two conditions, as localist and distributed strategies should both be available to solve this task. We consider the speeded recognition task to be an implicit assessment of AC knowledge because participants were not instructed to find AC associations, and the explicit inference task to be explicit in the sense that finding indirect pairs was instructed. This distinction mirrors that in the memory literature between slower associative retrieval-based processes versus more automatic recognition (Nobel & Shiffrin 2001; Cohn & Moscovitch 2007). However, we do not assume that the two tasks necessarily tap into implicit vs. explicit learning processes or knowledge representations (Shanks & John 1994).

**Figure 1.**
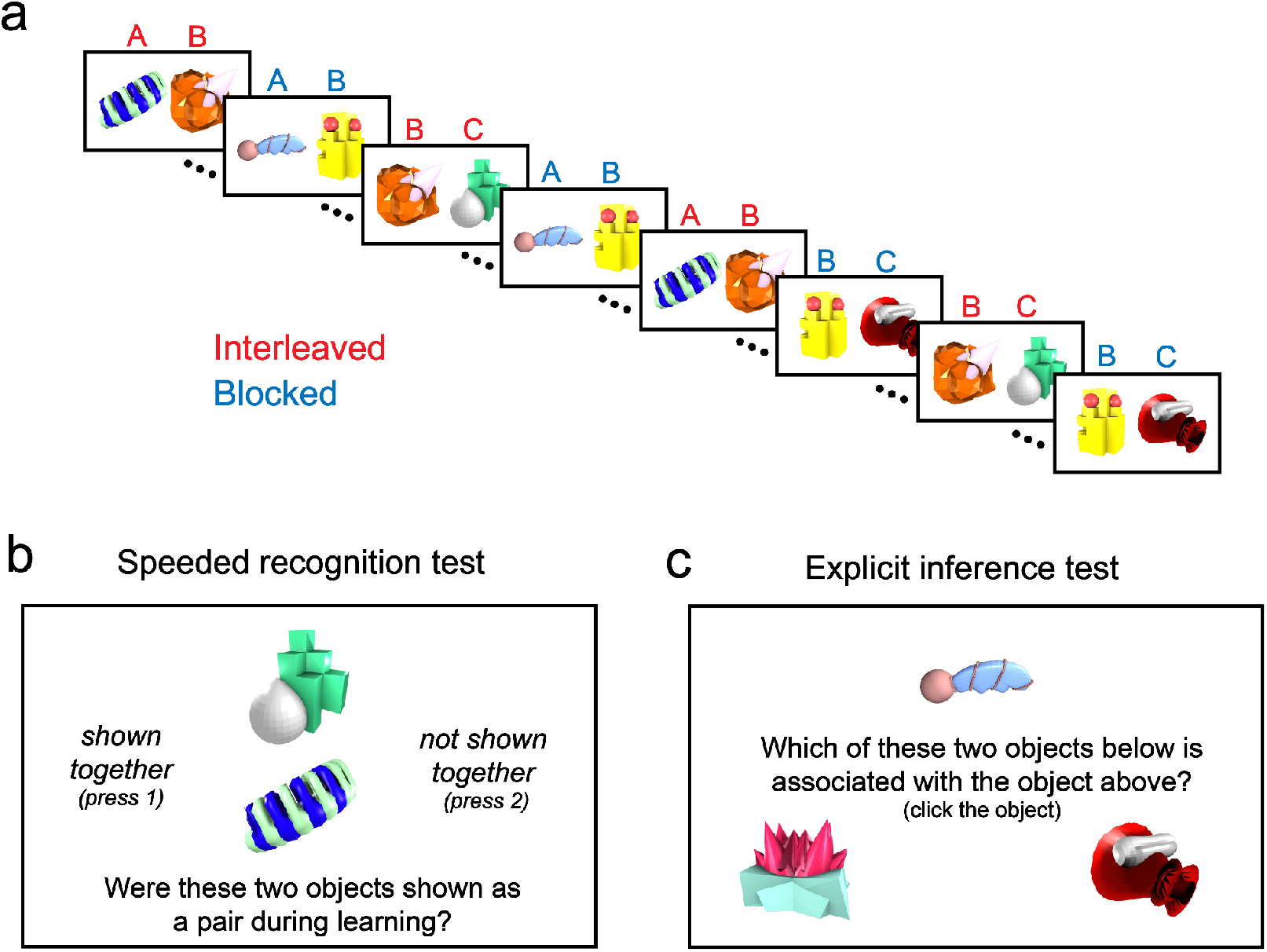
Exps 1a-c design. (a) During learning, participants viewed pairs of objects. Pairs that shared a common object (i.e., AB and BC) appeared either in interleaved (red) or blocked (blue) order. The relative horizontal order of items within each pair was randomized (not shown here). (b) In the speeded recognition test, participants quickly judged whether two objects were paired during learning. (c) In the explicit inference test, participants selected which of two objects was indirectly related to a cue object.

### Methods

#### Participants

In Experiment 1a, we recruited 33 participants, with 26 participants remaining after exclusions (7 females, 18 males, 1 unknown; 19 White, 2 Black, 3 Asian, and 2 unknown; 1 Hispanic or Latino; mean age = 33.24, SD = 7.70). Experiment 1b was a pre-registered study (https://osf.io/ag42z) in which 83 new participants were recruited, with 50 participants after exclusions (21 females, 29 males; 33 White, 5 Black, 9 Asian, and 3 unknown; 4 Hispanic or Latino; mean age = 38.08, SD = 9.07). The sample size for Experiment 1b was determined by a power analysis with a target power of 0.95 based on the effect size of the primary effect (i.e., difference in reaction time between speeded interleaved and blocked AC trials) observed in Experiment 1a. Experiment 1c was an additional replication study in which we recruited a total of 184 new participants, with 96 participants after exclusions (51 females, 44 males, 1 non-binary; 74 White, 8 Black, 1 American Indian/Native, 11 Asian, and 2 unknown; 5 Hispanic or Latino; mean age = 36.41, SD = 9.91). We collected a larger sample size in Experiment 1c for the purposes of analyses unrelated to this paper. Across all experiments, participants were recruited through Amazon Mechanical Turk (MTurk). Participants indicated their gender in a text box and reported ethnicity by selecting from “Hispanic or Latino”, “Not Hispanic or Latino”, and “Prefer not to answer”. Participants indicated their race by selecting from “American Indian or Alaska Native”, “Asian”, “Black or African American”, “Native Hawaiian or Other Pacific Islander”, “White”, “More than one race”, “Unknown”, and “Prefer not to answer”. The study protocol was approved by the local Institutional Review Board.

Across all three experiments, we excluded participants with d-prime lower than 1.5 on the speeded recognition task. Our primary motivation for this criterion was to exclude participants who showed poor memory for the studied direct pairs, since our theoretical predictions regarding AC inference assume knowledge of direct pairs, and prior associative inference studies have required and demonstrated strong memory for direct pairs (Koster et al., 2018; Shohamy & Wagner 2008; Schlichting et al., 2015). This exclusion criterion departed from our pre-registered plan, which only considered the hit rate in the speeded task, as we realized that it is also important to identify participants with high false alarm rates (participants who failed to reject objects that were neither directly nor indirectly related). Based on this exclusion criterion, we excluded 7 out of 33 (or 21.2%) participants in Exp 1a, 33 out of 83 (or 39.8%) participants in Exp 1b, and 88 out of 184 (or 47.8%) participants in Exp 1c. Our exclusion rates are high but comparable to the proportions of excluded participants often reported in studies conducted on online platforms such as Amazon Mechanical Turk and Prolific (e.g., Ludwin-Peery et al., 2021; Luo & Zhao 2018; Downs et al., 2010; Bokeria & Henson 2022; Himberger et al., 2019). While it was important, given our theoretical predictions, to limit our analyses to participants who demonstrated memory for direct pairs, we note that these exclusions may limit the generalizability of our findings.

#### Design and procedure

We adopted a within-subject design in which, during learning, each participant was shown a sequence consisting of presentations of 12 pairs of novel visual objects. These pairs were made up of a total of 18 items randomly sampled, for each participant, from 36 artificial object images (Hsu et al., 2014; Schlichting et al., 2015). Each “direct” pair AB was uniquely related to another direct pair BC through a linking item B. Each direct pair was shown 30 times. Among the six ABC triads shown to each participant, three were interleaved, with AB and BC appearing in alternation, and three were blocked, with all presentations of AB occurring before the first presentation of BC (Fig. 1a). Presentation of interleaved and blocked pairs was intermixed throughout the learning phase, and there was no demarcation of the halfway point. Pairs that share an object were never shown back-to-back. Participants were instructed to remember the pairings of objects by creating quick narratives of how the two objects might interact. Each participant completed a total of 360 trials, with the two objects displayed side-by-side on the screen for 1000 ms on each trial. Across repeated presentations of each object pair, the horizontal position of one item relative to the other was randomized (not counterbalanced). Subsequent to each pair presentation, participants saw the question “on a scale of 1 (failed to visualize a story) to 5 (clearly visualized a story), how well were you able to visualize a story linking the objects?” Participants made a total of 30 ratings for each pair. Participants responded with a numerical rating with a 7000 ms response deadline in Exps 1a and 1b, and a 2000 ms response deadline in 1c.

After learning, participants completed two tasks that probed their memory of learned object associations: a speeded recognition task (Fig. 1b) followed by an explicit inference task (Fig. 1c). During the speeded recognition task, on each trial, two objects were displayed for 1500 ms and participants were asked to respond within 3500 ms whether the two objects had been shown as a pair during the learning phase. The task comprised 24 trials with directly paired objects (e.g., AB), 12 trials with indirectly related objects (AC), and 12 trials with unrelated foil pairs (e.g., AD). Each object pair appeared in two trials, with two different vertical positions (A above C or C above A), with the constraint that pairs could not be repeated on back-to-back trials. At the onset of the explicit inference task, participants were instructed that two items paired with the same item form an indirect association (i.e., A and C items paired with the same B) and that they had to identify indirectly related items in this task. On each trial, participants saw a cue object at the top of the screen (A or C) and were instructed to select which of two objects shown below the cue object was indirectly related to it. The foil was of the same type as the target (and from the same interleaved or blocked condition) but from a different triad. There was a 7000 ms response deadline. The explicit inference task consisted of 12 trials where each AC association was tested twice, such that each item served as the cue object in one trial and as the target object in the other.

In Exp 1c, participants then completed an additional task that employed the same design as the explicit inference task, assessing memory for direct AB and BC pairs. This task consisted of 24 trials: 12 AB and BC pairs each shown twice, with each object shown as the cue in one trial and the target in the other. On each trial, participants selected between a target object and a matched foil with a 7000 ms response deadline.

#### Analysis

For each reaction time (RT) analysis, we excluded participants who had no data recorded for a condition of interest (e.g., missed all interleaved AC trials for the analysis of interleaved – blocked RT difference). Across all experiments, RTs for trials during which participants responded correctly were log-transformed before averaging. For each variable of interest, we performed paired two-tailed t-tests to assess the significance of within-subject differences between two conditions of interest. We additionally computed Bayes factors using the Jeffreys-Zellner-Siow prior as in Rouder et al. (2009) to quantify the relative likelihood between the hypothesis *H*_1_ of a difference between conditions and the null hypothesis *H*_0_ that no difference exists between conditions given the data. *H*_0_is favored when *B*_01_is greater than 1 whereas *H*_1_ is favored when *B*_1 0_ is greater than 1.

#### Transparency and openness

We report sample size, data exclusion, and details of methods, materials, design, and analysis in accordance with journal article reporting standards (Kazak 2018). All data and analysis code are available at https://github.com/schapirolab/itlblklearning. We pre-registered the design and analysis of Experiment 1 on OSF (https://osf.io/ag42z/) and performed replications for Experiments 1 and 3.

### Results and Discussion

During learning, participants responded to 97.96% of trials in Exp 1a (SE=1.8), with mean visualization score of 3.80 (SE=0.16), 99.03% of trials in Exp 1b (SE=0.8), with mean visualization score of 3.54 (SE=0.13), and 90.74% of trials in Exp 1c (SE=2.6) with mean visualization score of 3.37 (SE=0.10). The overall high response rates indicate that participants were attentive in the learning phase. Lower response rates in Exp 1c than in Exps 1a-b were likely due to the shorter response window. In the speeded recognition task, on trials where a response was made, participants indicated that they recognized the object pair 50.17% of the time in Exp 1a (SE=0.99), 51.87% of the time in Exp 1b (SE=0.78), and 52.56% of the time in Exp 1c (SE=0.76). (The correct proportion is 50%.)

In the speeded recognition task in Exp 1a, consistent with our predictions, participants were slower to correctly reject ACs as not having been paired during learning in the interleaved relative to blocked condition (Fig. 2a): On correct trials, participants responded slower to interleaved Acs than to blocked ACs (mean interleaved?blocked RT=0.061, SE=0.025, t(25)=2.41, p=0.02, Cohen’s d=0.47, *B*_10_=2.32; Fig. 2a) and to interleaved foils (i.e., pairs of objects from different interleaved triads; mean difference=0.13, SE=0.023, t(25)=5.56, p<0.001, d=1.09, *B*_10_=2413.72; Fig. S1), whereas RT did not significantly differ between blocked ACs and foils (mean difference=0.017, SE=0.018, t(25)=0.94, p=0.35, d=0.19, *B*_01_=3.23; Fig. S1) nor between interleaved and blocked foils (mean difference=-0.05, SE=0.028, t(25)=-1.78, p=0.09, d=0.34, *B*_01_=1.28; Fig. S2). RT did not differ between interleaved and blocked direct pairs (mean difference=-0.030, SE=0.022, t(25)=-1.38, p=0.18, d=0.27, *B*_01_=2.08; Fig. S10).

**Figure 2.**
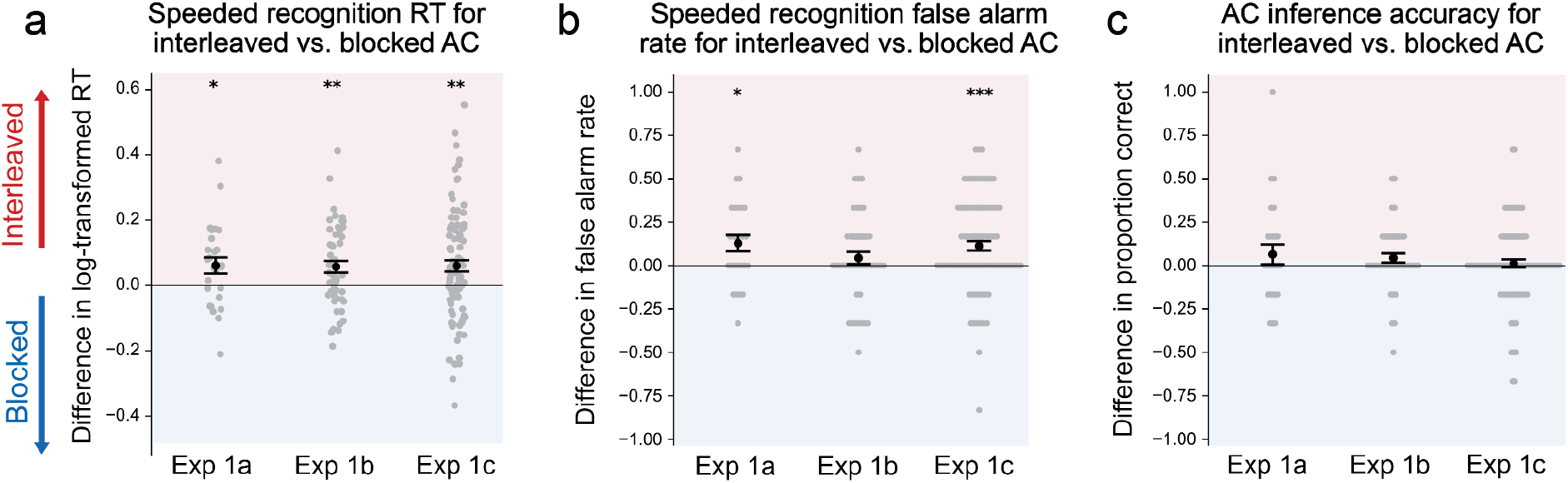
Exps 1a-c results. (a) RTs for interleaved vs. blocked AC trials during speeded recognition. (b) False alarm rates for interleaved vs. blocked AC trials during speeded recognition. (c) Accuracy differences between interleaved and blocked AC trials during explicit inference. Gray dots correspond to individual participants. Error bars represent +/-1 SEM. *p<0.05; **p<0.01; ***p<0.001.

In addition to our main prediction of an increased response time, we predicted that interleaving may also produce higher false alarm rates. Indeed, participants exhibited higher false alarm rates to interleaved than to blocked ACs (mean difference=0.13, SE=0.047, t(25)=2.70, p=0.01, d=0.53, *B*_10_=4.01; Fig. 2b), meaning they were more likely to incorrectly indicate that they had seen these indirect pairs when the direct constituents were presented in interleaved order. There was no difference in false alarm rates between interleaved and blocked foils (mean difference=0.013, SE=0.026, t(25)=0.49, p=0.63, d=0.10, *B*_01_=4.32; Fig. S3). These results support our prediction of an advantage of interleaved exposure for forming representations that support direct and automatic AC association.

To increase confidence in the results from Exp 1a, we preregistered (https://osf.io/ag42z/) and ran a replication of the experiment as Exp 1b. In the preregistration of Exp 1b, we indicated that our main prediction was increased response time on interleaved AC trials, but we also hypothesized increased false alarms to interleaved ACs in the speeded task. Exp 1b confirmed our main prediction: participants again responded significantly more slowly to interleaved ACs than to blocked ACs (mean=0.057, SE=0.018, t(49)=3.15, p=0.003, d=0.44, *B*_10_=11.35; Fig. 2a) and to interleaved foils (mean=0.10, SE=0.024, t(49)=4.37, p<0.001, d=0.62, *B*_10_=342.91; Fig. S1), and RT was not significantly different between blocked ACs and blocked foils (mean=0.041, SE=0.021, t(49)=1.94, p=0.06, d=0.27, *B*_01_=1.16; Fig. S1) or between interleaved and blocked foils (mean=-0.0043, SE=0.019, t(49)=-0.23, p=0.82, d=0.032, *B*_01_=6.34; Fig. S2). There was again no difference in RT between interleaved and blocked direct pairs (mean=-0.03, SE=0.017, t(49)=-1.80, p=0.078, d=0.25, *B*_01_=1.46; Fig. S11). Though numerically in the same direction as Exp 1a, we did not observe a significant difference in false alarm rates between interleaved and blocked AC trials in Exp 1b (mean=0.043, SE=0.035, t(49)=1.23, p=0.22, d=0.17, *B*_01_=3.20; Fig. 2b).

Exps 1a and b provide evidence for our main prediction of an interleaved advantage for rapid AC association. However, Exps 1a and b only measured memory for AB and BC pairs in the speeded recognition task, which does not assess the ability to explicitly identify direct associations against competing alternatives. To verify that participants can do this equally well for the two conditions, we performed an additional replication study, Exp 1c, which additionally assessed explicit identification of AB/BC pairs against foils at the end of the experiment. Results of Exp 1c again demonstrated stronger AC associations in the interleaved condition: During the speeded recognition task, participants responded more slowly to interleaved ACs than to blocked ACs (mean=0.060, SE=0.017, t(93)=3.41, p=0.001, d=0.35, *B*_10_=23.54; Fig. 2a) and to matched foils (mean=0.093, SE=0.018, t(94)=5.20, p<0.001, d=0.53, *B*_10_=11793.94; Fig. S1). RT again did not significantly differ between interleaved and blocked direct pairs (mean=0.0039, SE=0.011, t(95)=0.37, p=0.72, d=0.037, *B*_01_=8.30; Fig. S12). In Exp 1c we again observed, as in Exp 1a, higher false alarm rates to interleaved than to blocked ACs (mean=0.11, SE=0.027, t(95)=4.13, p<0.001, d=0.42, *B*_10_=235.86; Fig. 2b). RT (mean=0.024, SE=0.015, t(95)=1.64, p=0.10, d=0.17, *B*_01_=2.42; Fig. S2) and false alarm rates (mean=0.028, SE=0.018, t(95)=1.54, p=0.13, d=0.16, *B*_01_=2.85; Fig. S3) were not significantly different between interleaved and blocked foils. The strong false alarm effect arising again in this highly powered replication suggests that the lack of effect in Exp 1b may have been spurious.

In the task that explicitly assessed AB/BC memory, participants showed matched memory between the two conditions (mean=-0.0061, SE=0.0093, t(95)=-0.66, p=0.51, d=0.067, *B*_01_=7.19; Fig. S4a) and between AB and BC pairs in both conditions (interleaved: mean=0.012, SE=0.012, t(95)=1.04, p=0.30, d=0.11, *B*_01_=5.23; blocked: mean=-0.014, SE=0.0095, t(95)=-1.47, p=0.15, d=0.15, *B*_01_=3.14; Fig. S15). This result combined with the lack of differences in direct pair RTs suggest that the observed advantage for rapid recognition of interleaved AC associations is unlikely to be due to a difference in direct pair memory between conditions.

We examined whether recognition of individual direct pairs predicted performance on related indirect pairs in the speeded task in Exp 1c. As direct pair recognition accuracy was perfect for many participants, we conducted the analysis using RT. We did a median split for each participant on direct pair RT, and assessed whether responses to ACs were faster when the constituent pairs were faster. We observed no evidence of such a difference across different types of direct trials (interleaved AB: t(86)=-0.70, p=0.49, d=0.075, *B*_01_=6.68; interleaved BC: t(85)=0.42, p=0.67, d=0.046, *B*_01_=7.70; interleaved AB and BC: t(82)=-0.82, p=0.42, d=0.09, *B*_01_=5.98; blocked AB: t(81)=-0.60, p=0.55, d=0.066, *B*_01_=6.91; blocked BC: t(83)=0.75, p=0.46, d=0.08, *B*_01_=6.35; blocked AB and BC: t(76)=-0.52, p=0.60, d=0.060, *B*_01_=6.98), suggesting that response speed to direct pairs does not predict response speed to their indirect associations in this study.

Participants responded slower during the explicit inference task than in this explicit direct pair assessment in both the interleaved (mean=0.38, SE=0.022, t(95)=17.10, p<0.001, d=1.74, *B*_10_>1.0e+6; Fig. S9) and the blocked condition (mean=0.40, SE=0.024, t(94)=16.60, p<0.001 d=1.70,, *B*_10_>1.0e+6). We also observed slower responses to indirect than to direct speeded recognition trials across Exps 1a (interleaved: mean=0.21, SE=0.027, t(25)=7.73, p<0.001, d=1.52, *B*_10_=324731.5; blocked: mean=0.12, SE=0.022, t(25)=5.18, p<0.001, d=1.02, *B*_10_=988.07; Fig. S8), 1b (interleaved: mean=0.21, SE=0.021, t(49)=10.332, p<0.001, d=1.46, *B*_10_>1.0e+6; blocked: mean=0.13, SE=0.023, t(49)=5.51, p<0.001, d=0.78, *B*_10_=12552.24), and 1c (interleaved: mean=0.18, SE=0.019, t(94)=9.3625, p<0.001, d=0.96, *B*_10_>1.0e+6; blocked: mean=0.12, SE=0.014, t(94)=8.6209, p<0.001, d=0.88, *B*_10_>1.0e+6), suggesting that direct associations were always more accessible than indirect associations. Exps 1a, 1b, and 1c together provide strong evidence that interleaved exposure facilitates direct and automatic association of indirectly related items.

As predicted, across Exps 1a, 1b, and 1c, we did not observe significant accuracy differences between inference performance for interleaved and blocked ACs in the explicit inference task (1a: mean=0.064, SE=0.059, t(25)=1.10, p=0.28, d=0.21, *B*_01_=2.81; 1b: mean=0.043, SE=0.027, t(49)=1.61, p=0.11, d=0.23, *B*_01_=1.94; 1c: mean=0.012, SE=0.024, t(95)=0.51, p=0.61, d=0.052, *B*_01_=7.82; Fig. 2c, Fig. S16). The *B*_01_value of 7.82 for Exp 1c, which had the highest power of the three versions of Exp 1 to detect an effect, provides especially strong evidence for a lack of difference between conditions. There were also no differences in RT (1a: mean=-0.030, SE=0.066, t(24)=-0.45, p=0.66, d=0.090, *B*_01_=4.32, Fig. S10; 1b: mean=0.003, SE=0.038, t(49)=0.085, p=0.93, d=0.012, *B*_01_=6.48, Fig. S11; 1c: mean=-0.020, SE=0.023, t(94)=-0.86, p=0.39, d=0.088, *B*_01_=6.18, Fig. S12). Equivalent performance between the two conditions in this relatively slow and explicit setting indicates that participants were able to successfully link blocked AC associations despite displaying little sensitivity to these associations in the implicit speeded setting. This pattern of results is consistent with the use of relatively slower recurrent computation across localist representations, as hypothesized by REMERGE (Kumaran & McClelland 2012; Tamminen et al., 2015).

## Experiment 2

Exps 1a-c demonstrated an enhanced sensitivity to indirect associations after interleaved learning, but since the task asked participants to report only the directly learned associations, interleaving was arguably not advantageous in that setting. In the next set of experiments, we aimed to highlight situations in which representations learned through interleaving should be beneficial. In neural network models, distributed representations support automatic generalization (Hinton, 1984): Learning new knowledge about an object automatically allows that knowledge to be transferred to related objects by virtue of the overlap in their representations. By contrast, localist codes do not as naturally support such transfer (they can, but only with additional carefully constructed recurrent computation; McClelland et al., 1986). In Exp 2, we tested whether behavior after interleaved learning reflects this quality of distributed representations. After participants learned interleaved and blocked AB and BC associations as in Exp 1 (Fig. 3a), they then learned to associate novel nonsense properties with a subset of the objects (Fig. 3b). For each ABC triad, participants associated a unique property with either A or C. Participants then completed three tests: property memory, property generalization, and explicit direct pair recognition. In the property memory test (Fig. 3c), we assessed participants’ memory for the trained property-object pairings, with no expectation of a difference in performance between conditions. In the property generalization test, we assessed their ability to generalize the novel properties to indirectly related objects (Fig. 3d). We did not explicitly instruct them about the presence of indirectly related objects, as we had in the explicit inference task above. We predicted superior property generalization for interleaved relative to blocked ACs. Finally, participants were explicitly assessed on their memory for AB and BC pairings as in Exp 1c (not shown in Fig. 3).

**Figure 3.**
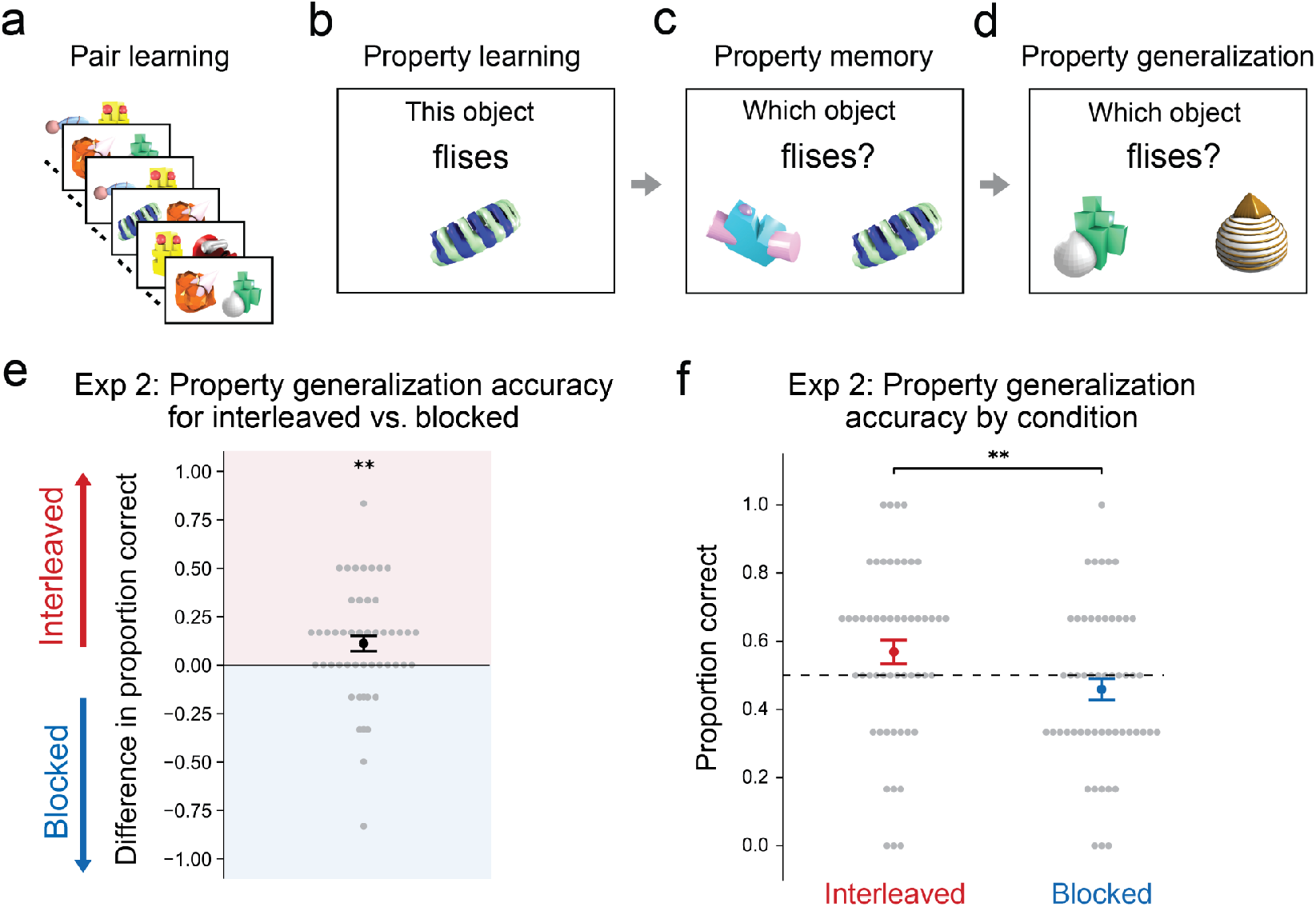
Exp 2 design and results. (a) Participants viewed ABC triads in either interleaved or blocked order as in Exp 1. (b) During property learning, participants learned to associate a subset of objects with novel nonsense properties. (c) The property memory test assessed participants’ memory for these trained pairings. (d) In the property generalization task, participants were asked to identify which of two objects was more likely to have the novel property, one of which was indirectly related to an object that participants had learned had that property. (e) Difference in property generalization accuracy between interleaved and blocked triads. (f) Property generalization accuracy broken down by condition. Dashed line indicates chance performance.

### Methods

#### Participants

In Experiment 2, to match the sample size of the pre-registered Experiment 1b, we recruited 84 new participants through MTurk. We excluded participants who missed more than half of explicit direct pair memory trials or more than one third of property memory trials, resulting in a total of 53 participants (25 females, 28 males; 42 White, 5 Black, 2 Asian, and 4 unknown; 5 Hispanic or Latino; mean age = 37.66, SD = 11.87), which corresponds to an exclusion rate of 36.9%. We chose these criteria to exclude participants who showed weak direct pair memory or property memory while retaining enough data for our analyses.

#### Design and procedure

After learning direct pair associations as in Exps 1a-c, participants learned novel properties of a subset of the objects, and then completed a property memory and property generalization task. During property learning (Fig. 3b), each participant either learned to associate the six A objects (26 participants) or the six C objects (27 participants) across both conditions with six novel nonwords, including “smobs”, “cwoads”, “flises”, “loarts”, “misks”, and “jolmbs”, selected from the ARC nonword database (Rastle et al., 2002). Prior to property learning, we instructed participants that each of the objects they saw had a hidden property and that they would learn some of these properties. On each property learning trial, an object was displayed for 2500 ms along with a text description of its associated property (“this object flises”). After each object-nonword association presentation, participants rated on a scale of 1 to 5 how well they felt they were able to learn the association, with a 4000 ms response deadline. Each object-property association was displayed 12 times during property learning.

During the property memory task (Fig. 3c), on each trial, participants saw a property at the top of the screen (e.g., “flises”) and were instructed to select, between a target object and a matched foil, the object that had been associated with the property during learning. There was a 4000 ms response deadline. Each property association was tested twice, with two different horizontal positions (i.e., target on the left and foil on the right, and vice versa), for a total of 12 trials.

Before the property generalization task (Fig. 3d), participants were instructed that they would see a property they had studied along with two objects that they had not learned hidden properties for, and they would be asked to judge which of the two objects also has that property. On each trial, participants saw a property at the top of the screen (e.g., “flises”) and selected between a target object and a matched foil, with a 4000 ms response deadline. The target object was always indirectly related to the associated object (i.e. A to C). Each object-nonword association was tested twice, with two different horizontal positions (i.e., target on the left and foil on the right, and vice versa), forming a total of 12 trials. Finally, participants were explicitly assessed on their memory for AB and BC pairings as in Exp 1c.

### Results and Discussion

In Exp 2, during learning, participants responded to 93.02% of all trials (SE=2.72) with a mean visualization score of 3.40 (SE=0.14). As expected, participants’ memory for trained object-property associations was not different between interleaved (mean=0.93, SE=0.016) and blocked (mean=0.93, SE=0.019) items in the property memory test (mean difference=0.0, SE=0.02, t(52)=-0.00, p=1.00, d=0.0, *B*_01_=6.68; Fig. S6). However, in the property generalization task, participants performed significantly better for interleaved (mean=0.57, SE=0.035, t(52)=1.98, p=0.053, d=0.27, *B*_01_=1.10; Fig. 3f) than blocked (mean=0.46, SE=0.032, t(52)=-1.29, p=0.20, d=0.18, *B*_01_=3.06; Fig. 3f) triads (mean difference=0.11, SE=0.040, t(52)=2.78, p=0.008, d=0.38, *B*_10_=4.65; Fig. 3e).

Unlike Exp 1c, participants here showed superior explicit memory (mean difference=0.035, SE=0.017, t(52)=2.07, p=0.04, d=0.28, *B*_10_=1.10; Fig. S5a) for interleaved (mean=0.86, SE=0.022; Fig. S5a) relative to blocked direct pairs (mean=0.83, SE=0.021; Fig. S5a). This raises the possibility that weak property generalization performance in the blocked condition could be due to poorer memory of blocked direct pairs, though there was no reliable correlation between subjects’ overall accuracy on blocked pairs and performance on property generalization (r=0.021, p=0.83). To rule out this possibility, we ran analyses restricted to a subsample of subjects (n=44) who had matched performance between conditions on direct pairs (mean=0.0019, SE=0.016, t(43)=0.12, p=0.91, d=0.02, *B*_01_=6.08), by excluding subjects who showed the weakest memory for blocked relative to interleaved pairs. Analyses based on this sample again revealed matched memory for object-property associations between conditions (mean=0.0038, SE=0.025, t(43)=0.15, p=0.88, d=0.023, *B*_01_=6.06) and superior property generalization performance for interleaved relative to blocked triads (mean=0.10, SE=0.044, t(43)=2.34, p=0.024, d=0.35, *B*_10_=1.87). Thus, the advantage of interleaved associations during property generalization is unlikely to be due to lack of retention of direct pairs in the blocked condition. In sum, Exp 2 provides evidence that representations formed via interleaved exposure afford automatic generalization of new knowledge to indirectly related objects.

## Experiments 3a-b

Thus far, we have demonstrated that interleaved learning influences behavior in tasks that require relatively rapid judgments. Exps 1a-c did not identify a difference between interleaved and blocked conditions in the slower, explicit AC inference task. Indeed, we did not predict such a difference, due to the availability of an alternative strategy in this case: indirect association through recurrent computation across localist representations, as proposed by the REMERGE model (Kumaran & McClelland, 2012). Are there certain learning problems where, even given sufficient opportunity for retrieval-based computation, only interleaved exposure can support successful behavior? One situation in which we expect the localist strategy to fail is when direct associations are not clearly demarcated during encoding, as AC inference using the localist strategy requires clean conjunctive representations of direct AB and BC pairs (Kumaran & McClelland, 2012; Schapiro et al. 2017). In temporal statistical learning paradigms, objects are presented one at a time in a continuous stream with embedded regularities (Saffran et al., 1996). A localist strategy that quickly forms robust conjunctive representations of every observed temporal co-occurrence would encode both reliable and unreliable pairings. AC inference using this strategy would be very difficult, as recurrent processing at retrieval would activate the unreliable associations. In contrast, distributed representations are very well suited to capturing graded statistical regularities (Rogers & McClelland 2004; Schapiro et al. 2017). We therefore predicted that a statistical learning variant of our paradigm would show a robust advantage for interleaved exposure even in the slow explicit inference test.

### Methods

#### Participants

To determine the sample size required to detect reliable effects in Experiment 3, we found a prior visual object statistical learning study that evaluated recognition of object pairs (Park et al. 2018). This study had samples of 30 participants in each experiment, and an average effect size *d* of 0.77 for performance on direct pairs, which corresponds to a sample size of 24 to achieve 0.95 power. We aimed to collect at least 30 participants after exclusions for these studies (with variance above this due to uncertainty in yield with online batch data collection). In Experiment 3a, 105 new participants were recruited through MTurk. To exclude participants who showed inattentiveness during learning or weak knowledge of direct associations after learning, we excluded participants who missed more than half of learning phase responses or more than one third of speeded direct trials, resulting in 43 participants (17 female, 26 males; 37 White, 1 Black, 1 American Indian/Native, 3 Asian, and 1 unknown; 4 Hispanic or Latino; mean age = 37.49, SD = 11.61). Experiment 3b was a replication study in which 100 new participants were recruited through MTurk, resulting in 35 participants after exclusions (15 female, 19 males, 1 unknown; 26 White, 6 Black, 2 Asian, and 1 unknown; 3 Hispanic or Latino, 1 unknown; mean age = 34.71, SD = 10.59). The exclusion rates were 58.1% and 65% in Exp 3a and 3b, respectively, which are higher than those in previous experiments, reflecting the fact that learning direct associations is more challenging in the statistical learning setting. Indeed, it can be difficult to identify evidence of successful statistical learning in online experiments (Himberger et al., 2019). Because our hypotheses regarding AC inference are motivated by models that assume knowledge of the direct associations, it was important to restrict analyses to participants who successfully learned direct associations, but, as before, this approach may limit the generalizability of our results.

#### Design and procedure

For each participant, a sequence of visual object pairs was generated following the same protocol as Exps 1a-c, except that each pair repeated 24 (instead of 30) times, and objects were presented one at a time in a continuous stream with no breaks, such that two objects from the same pair were shown consecutively followed by objects from a different pair (Fig. 4a). For each occurrence of an object pair, the order of the objects was randomized. Therefore, for an object pair AB or BC, the two objects always appeared adjacent in the sequence, but each object would also appear adjacent to objects that were neither directly nor indirectly related. As a cover task, participants were instructed to quickly respond as to whether the current object appeared heavier than the previous object in the sequence. Participants pressed one key if the current object seemed heavier than the preceding object, and a different key if not. Each object was displayed for 1000 ms followed by an intertrial interval of 500 ms. After learning, participants were informed that there were object pairs embedded within the sequence they saw. Participants then completed the speeded recognition task followed by the explicit inference task as in Exps 1a-c. In 3b, participants additionally completed an explicit direct pair recognition task as in Exps 1c and 2.

**Figure 4.**
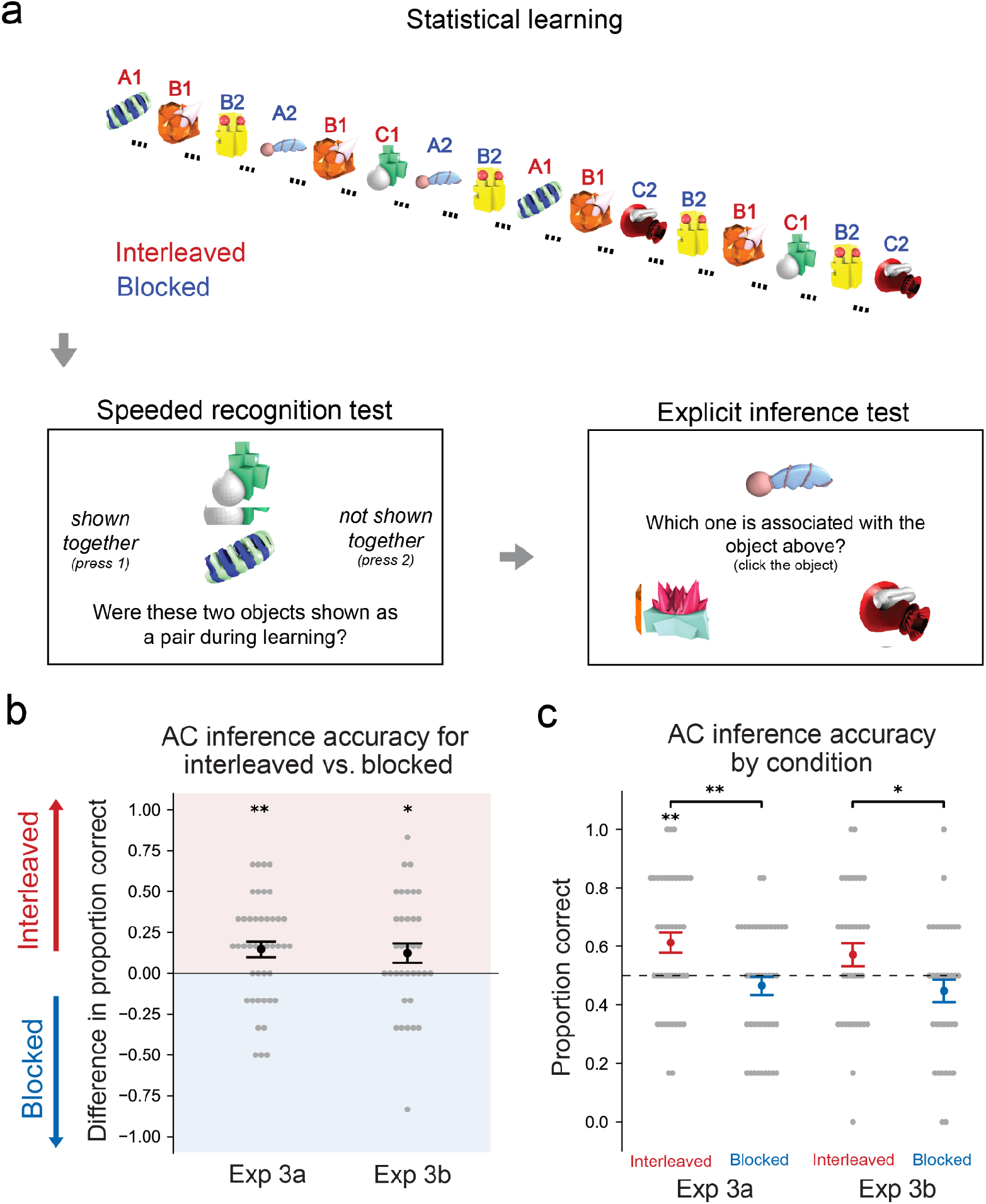
Exps 3a-b design and results. (a) Participants viewed objects presented one at a time followed by a speeded recognition test and explicit inference test. (b) Explicit inference accuracy differences between interleaved and blocked trials in Exp 3a and 3b. (c) Explicit inference accuracy by condition in Exp 3a and 3b.

### Results and Discussion

In Exp 3a, in the learning phase, participants responded to an average of 93.78% of all trials (SE=1.55) and indicated that an object appeared heavier than its preceding object on 51.48% of the trials they responded to (SE=1.28). In Exp 3b, participants responded to an average of 91.36% learning trials (SE=1.50) and indicated that an object appeared heavier than its preceding object on 53.03% of the learning trials they responded to (SE=1.10). These numbers suggest that participants were attentive during learning. As in Exps 1a-c, in the speeded recognition task, 50% of trials contained pairs that were actually shown during learning. On trials where a response was made, participants indicated that they saw an object pair during learning 64.21% of the time in Exp 3a (SE=1.59) and 69.64% of the time in Exp 3b (SE=2.05). These high rates relative to prior studies indicate that statistical learning induces higher familiarity for some unpaired objects (as discussed below).

As in Exps 1a-c, we observed evidence of stronger rapid recognition of interleaved relative to blocked indirect associations. Participants were again slower to correctly reject interleaved relative to blocked ACs in 3a (mean difference=0.081, SE=0.032, t(37)=2.50, p=0.017, d=0.41, *B*_10_=2.66; Fig. S7b) though not in 3b (mean=-0.013, SE=0.06, t(25)=-0.21, p=0.83, d=0.041, *B*_01_=4.73; Fig. S7b), whereas there was no difference between interleaved and blocked foils in 3a (mean=-0.021, SE=0.028, t(36)=-0.74, p=0.47, d=0.12, *B*_01_=4.40; Fig. S7b) or 3b (mean=-0.0067, SE=0.043, t(27)=-0.16, p=0.88, d=0.03, *B*_01_=4.93; Fig. S7b). Participants exhibited higher false alarm rates for interleaved than blocked ACs in both 3a (mean=0.26, SE=0.047, t(42)=5.42, p<0.001, d=0.83, *B*_10_=6726.25; Fig. S7a) and 3b (mean=0.19, SE=0.066, t(34)=2.81, p=0.008, d=0.47, *B*_10_=5.05; Fig. S7a). Unlike in Exps 1a-c, however, we observed evidence that interleaving induced a false sense of familiarity even for unrelated objects in the interleaved condition: response time was not significantly different between interleaved ACs and foils in 3a (mean=0.095, SE=0.05, t(34)=1.88, p=0.069, d=0.31, *B*_01_=1.15) or 3b (mean=-0.012, SE=0.068, t(26)=-0.17, p=0.87, d=0.033, *B*_01_=4.84), and participants showed higher false alarm rates for interleaved than for blocked foils in 3a (mean=0.21, SE=0.049, t(42)=4.34, p<0.001, d=0.66, *B*_10_=271.20; Fig. S7a) but not in 3b (mean=0.10, SE=0.067, t(34)=1.57, p=0.13, d=0.27, *B*_01_=1.80; Fig. S7a). These results suggest that, when direct pair associations are not clearly demarcated, temporal proximity may facilitate the learning of associations even for unrelated items.

Critically, in Exp 3a, unlike in Exps 1a-c, we observed higher accuracy for interleaved than blocked AC pairs in the explicit inference task (mean=0.15, SE=0.049, t(42)=3.02, p=0.004, d=0.46, *B*_10_=8.30; Fig. 4b, Fig. S16), despite no difference in direct pair accuracy (mean=0.027, SE=0.034, t(42)=0.80, p=0.43, d=0.12, *B*_01_=4.49; Fig. S19) or RT (mean=-0.024, SE=0.019, t(42)=-1.26, p=0.21, d=0.19, *B*_01_=2.91; Fig. S13) between conditions. We found significant above chance inference performance in the interleaved condition (mean interleaved=0.61, SE=0.035, t(42)=3.22, p=0.0025, d=0.49, *B*_10_=13.23) but chance performance in the blocked condition (mean blocked=0.47, SE=0.032, t(42)=-1.10, p=0.28, d=0.17, *B*_01_=3.44), suggesting that only interleaved exposure permitted successful inference (Fig. 4c). Exp 3b largely replicated these effects: performance in the explicit inference task (mean interleaved=0.57, SE=0.04; mean blocked=0.45, SE=0.039) was superior in the interleaved condition (mean difference=0.12, SE=0.060, t(34)=2.06, p=0.047, d=0.35, *B*_10_=1.18; Fig. 4b, Fig. S16) despite matched direct pair performance (speeded recognition accuracy: mean=0.026, SE=0.030, t(34)=0.88, p=0.38, d=0.15, *B*_01_=3.85, Fig. S19; speeded recognition RT: mean=0.023, SE=0.028, t(34)=0.80, p=0.423 d=0.14, *B*_01_=4.10, Fig. S14; slow recognition accuracy: mean=-0.033, SE=0.036, t(34)=-0.94, p=0.35, d=0.16, *B*_01_=3.67, Fig. S4b; slow recognition RT: mean=-0.013, SE=0.029, t(33)=-0.44, p=0.66, d=0.076, *B*_01_=4.96, Fig. S14), indicating that the advantage in explicit inference for interleaved

ACs was not due to a failure to learn direct AB and BC associations in the blocked condition (though note enhanced memory for blocked BC relative to AB pairs in this case: mean difference=0.10, SE=0.048, t(34)=2.17, p=0.037, d=0.37, *B*_10_=1.42; Fig. S15). As in Exp 1c, participants were slower during explicit recognition of indirect than direct pairs for both interleaved (mean=0.18, SE=0.038, t(33)=4.70, p<0.001, d=0.81, *B*_10_=525.20; Fig. S9) and blocked trials (mean=0.32, SE=0.068, t(32)=4.70, p<0.001, d=0.82, *B*_10_=507.02; Fig. S9). Together, the results indicate that in a setting where object pairings need to be inferred from the statistics of co-occurrence over time, interleaving benefits even explicit inference.

### Model simulations

To evaluate how different classes of models behave in our paradigm, we contrasted three models of memory that have been proposed to solve associative inference (Fig. 5): the temporal context model (TCM; Howard & Kahana 2002; Howard et al., 2009; Fig. 5a-c), which employs distributed representations, REMERGE (Kumaran & McClelland 2012; Fig. 5d-f), which uses only localist representations, and our model of the hippocampus, C-HORSE (based on Schapiro et al., 2017; Fig. 5g-i), which contains both kinds of representation separated across the two pathways of the hippocampus. Code for these models can be found at: https://github.com/schapirolab/itlblklearning.

**Figure 5.**
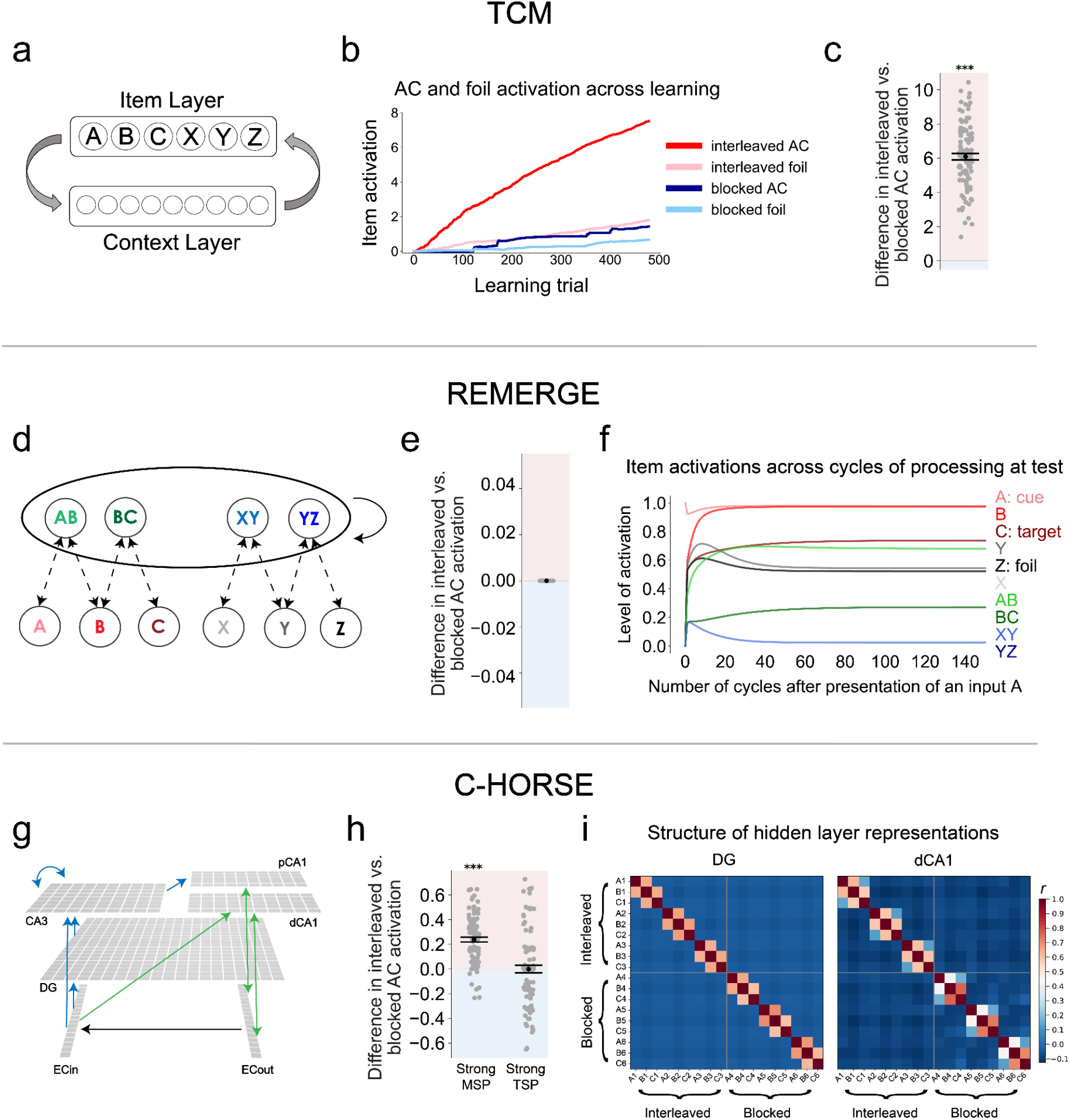
Model architectures and results. (a) TCM learns associations between an Item Layer and a Context Layer, which employs a distributed representation. (b) Throughout learning, indirectly related items A and C in the interleaved condition activate one another more than unrelated foils and those in the blocked condition. (c) Stronger activation of interleaved than blocked ACs at test across 100 simulations. (d) In REMERGE, AB and BC pairs are represented as idealized localist, conjunctive units connected to associated input units via bidirectional connections (dashed lines). (e) REMERGE predicts no difference between interleaved and blocked AC activation at test. (f) Item activations across a test trial after presentation of item A. As recurrent dynamics settle in the system, the activity of the target C is higher than that of matched foil Z. X is identical to Z and XY is identical to YZ. (g) C-HORSE is a neural network model instantiating known connectivity between subfields of the hippocampus. Blue connections make up the trisynaptic pathway (TSP) and green connections make up the monosynaptic pathway (MSP). (h) The model predicts an advantage for interleaved AC activation when behavior is driven predominantly by the MSP and equivalent AC activation between conditions when the TSP drives behavior. (i) Heatmap of hidden layer unit activity correlations in DG and dCA1 when individual items are presented. The “initial response” is used for these correlations, which is the pattern of activity evoked by the input before activity has time to spread from EC_out to EC_in on a particular trial (allowing cleaner assessment of the contributions of each region). DG shows similarity for directly (A1-B1) but not indirectly (A1-C1) related items, whereas dCA1 shows similarity for both directly and indirectly related items, especially in the interleaved condition.

### Overview of model methods

Here we provide an overview of the implementation and mechanisms of the three models we applied to our paradigm. Details and equations are provided in the Appendix. TCM encodes items by associating them with the contexts in which they are encountered. Relationships between items are then reflected in the degree of overlap in distributed context representations. TCM learns by updating item-to-context and context-to-item mappings. Prior to the experiment, distinct items are assumed to be associated with non-overlapping contexts. During learning, each input item causes drift in a context vector, and subsequent items are associated with this drifting context. Through this process, items’ associated contexts become overlapping to the extent that items continue to appear in similar contexts.

To simulate the task, we adopted a version of TCM designed to simulate paired associate learning (Howard et al., 2009; Supplementary Table 1). To delimit trials of pair presentations (as in Exp 1), a distractor vector drifts the context prior to each new trial. The context drift caused by an item is determined by the item representation and a recency-weighted average of past contexts of its occurrence. In the interleaved condition of our design, the context drift caused by each B will consistently reflect recent contexts in which its related A and C were presented. In contrast, the drift caused by each blocked item B will reflect only the contexts of its related A initially, until it becomes gradually swamped by that of its related C as BC pairs begin to appear. Prior context exponentially decays as new contextual information is incorporated. As a result of this interference, contexts in which interleaved A and C appear will be more similar than those of blocked A and C. Thus, in TCM, we expected that interleaved ACs would develop stronger associations than blocked ACs.

In contrast to the distributed representations used in TCM, REMERGE (Kumaran & McClelland 2012; Fig. 5d; Supplementary Table 2) employs a localist code: AB and BC are represented as non-overlapping units in the conjunctive layer, each of which is bidirectionally connected to individual item units in the feature layer (i.e., AB in the conjunctive layer is connected to A and B in the feature layer). This model employs an idealized learning process whereby each pair presentation strengthens connections only to constituent items, such that each AB presentation only updates bidirectional A-AB and B-AB connections. Since weight updates are fully orthogonalized for different conjunctive pairs, there is no basis for a difference between interleaving and blocking in the model’s behavior.

To infer item associations at retrieval, REMERGE engages in a search process by allowing activity to spread amongst connected units (McClelland et al., 1986). At the onset of retrieval, an external input activates the corresponding unit in the feature layer. At each timestep during retrieval, the net input into each unit is a weighted sum of its previous net input and the current inputs from connected units. In the feature layer, each unit’s activity is a logistic function of its net input. In the conjunctive layer, a version of the softmax function normalizes net inputs across units, mimicking competitive inhibition. AC inference can be achieved when an input A causes activity to spread to AB, then B, BC, and finally C. REMERGE is intended to simulate “big-loop” recurrence within the hippocampal system, in which output activity from the hippocampus can be recirculated back into the system as input via the entorhinal cortex.

C-HORSE (based on Schapiro et al., 2017; Fig. 5g), inspired by known circuitry and properties of the hippocampus, posits that both distributed and localist representations are available via separate pathways. The model consists of an input layer representing superficial layers of entorhinal cortex (EC_in), an output layer representing deep layers of entorhinal cortex (EC_out), and hidden layers representing hippocampal subfields dentate gyrus (DG), cornu ammonis 3 (CA3), and CA1. We modified the architecture of our previously published model (Schapiro et al., 2017; see Supplementary Tables 3-5 for comparison) to incorporate the topographic organization of projections from CA3 and superficial EC to CA1, along its proximodistal axis (Sun et al., 2014; Witter et al., 2006). We split the existing CA1 layer into two parts (<u>p</u>roximal/<u>d</u>istal CA1), such that the trisynaptic pathway (TSP) flow is EC_in -> DG -> CA3 -> pCA1 -> EC_out and the monosynaptic pathway (MSP) is EC <-> dCA1 (Fig. 5g). This allows the model to more easily express the distinct contributions of the MSP and TSP relative to our prior version. Otherwise, the model was the same as in Schapiro et al., 2017, with sparse connectivity and high inhibition in the TSP, which give rise to the well-known pattern-separated representations in DG and CA3 (Leutgeb et al., 2007), and dense connectivity and lower inhibition in the MSP, which produce relatively overlapping representations in dCA1. The TSP has a faster learning rate than the MSP, consistent with evidence suggesting that the TSP can do one-shot learning, whereas the MSP does more incremental learning (Nakashiba et al., 2008). We expected that overlapping representations in the MSP would require interleaving to support inference, whereas pattern-separated representations in the TSP would form via either interleaved or blocked exposure and support inference via recurrence, as in REMERGE.

For TCM and C-HORSE, we presented sequences of object pairs, in which AB and BC were either interleaved or blocked, generated using the same protocol and number of exposures as in Exps 1a-c. For REMERGE, we assumed that all connection weights are identical by the end of learning, as in its original simulations, where the extent of its weight strengthening depends only on the amount of training (i.e., the number of presentations), which is equivalent across all pairs. After learning, we measured the amount of activation of item C (as output) given A (as input), and vice versa, across 100 instantiations of each model.

### Results and Discussion

In TCM, as expected, interleaving led to stronger AC activation relative to blocking (t(99)=16.58, p<0.001, *B*_10_>1.0e+6) and relative to matched foil pairs (t(99)=13.29, p<0.001, *B*_10_>1.0e+6) across learning (Fig. 5b) and at test (Fig. 5c). In addition, there was greater activation for interleaved foils than blocked foils (at the end of training: t(99)=9.38, p<0.001, *B*_10_>1.0e+6; Fig. 5b), reflecting the fact that contexts associated with adjacent trials are not fully orthogonalized. TCM can thus account for situations where interleaving benefits inference, but cannot explain matched performance, as observed in the explicit inference task in Exp 1. We did not perform parameter fitting for this model, but found that the interleaved advantage was highly robust across parameter values, indicating a qualitative, inherent property of the model.

In the input sequences we generated, the delay between related pairs was, on average, shorter in the interleaved condition. Of the three models we considered, TCM is the only one that might plausibly be sensitive to this difference in time lag. To test whether the advantage of interleaving in TCM arises from this property of the input, we performed an additional simulation in which pairs from each triad in the blocked condition appeared only in the first half or in the second half of the full sequence. In this scenario, the mean time lag between related pairs is shorter in the blocked than in the interleaved condition, but we found that interleaving still resulted in stronger AC activation (Fig. S20) relative to blocking (relative to ACs blocked in the first half: t(99)=28.65, p<0.001, *B*_10_>1.0e+6; relative to ACs blocked in the second half: t(99)=29.17, p<0.001, *B*_10_>1.0e+6) and relative to matched foil pairs (relative to blocked foils in the first half: t(99)=29.1, p<0.001, *B*_10_>1.0e+6; relative to blocked foils in the second half: t(99)=29.0, p<0.001, *B*_10_>1.0e+6). Thus, associating related ACs in TCM benefits from the interleaved order of presentation, not simply the shorter time lags in the interleaved condition.

In REMERGE, given opportunities for spreading activation amongst related units, an input A always activates its associated items more than matched foils (Fig. 5f). This recurrent process first activates B and AB, more than foils Y and XY, followed by greater activation of BC than of YZ. Finally, C becomes activated more than its matched foil Z. The model predicts no difference between interleaved and blocked AC activations (Fig. 5e). This behavior is due to two critical features of REMERGE: there are no shared weights between conjunctive direct pairs, and learning depends on the number but not the order of presentations. REMERGE thus provides an account of how inference performance can be matched between conditions, but fails to explain the observed advantage of interleaved associations in the speeded recognition task.

It is worth considering whether modifications to REMERGE might allow it to show order effects that could explain the interleaving advantage. REMERGE could be modified to incorporate a weight decay mechanism, such that weights are higher for more recently presented item pairs. This would result in weaker memory for blocked ABs than BCs. Indeed, we generally see evidence for somewhat weaker AB relative to BC memory (Fig. S17). However, in many implementations of such a recency weighting, and, critically, in our data, memory for interleaved direct pairs is matched to memory for blocked direct pairs (Fig. S18), meaning that interleaved direct pair memory falls evenly between blocked AB and BC memory. REMERGE does not exhibit a stronger overall connection from A to C in the interleaved condition under the constraint of matched overall direct pair memory. It is possible that the matched memory we observed in Exp 1 was due to a ceiling effect, but recognition of direct pairs in Exp 3 was off ceiling and also matched between conditions (Fig. S19). Mismatched direct pair strength between conditions would also likely lead to differences in explicit inference performance in Exps 1a-c, which we did not observe. In sum, it is possible in theory that an implementation of recency weighting that results in stronger overall direct pair memory in the interleaved condition could lead to an interleaved advantage in speeded recognition for REMERGE, but we did not observe such an advantage in direct pair memory in our data.

In C-HORSE, we simulated the speeded and explicit inference tasks by modulating reliance on the MSP vs. TSP. We assumed a control mechanism that shifts the relative strength of dCA1/pCA1 outputs to EC_out during AC inference such that model dynamics were predominantly driven by only one of the two pathways at a time. Such a mechanism could potentially be implemented in interactions between medial prefrontal cortex and CA1, as medial prefrontal cortex is known to influence CA1 representations as a function of task requirements (Guise & Shapiro, 2017; Eichenbaum, 2017). We found that when relying primarily on the sparse, pattern-separated representations of the TSP (Fig. 5i), the model demonstrates matched performance between the interleaved and blocked conditions (t(99)=-0.03, p =0.98, *B*_01_=10.04; Fig. 5h), akin to human participants on the explicit inference task. On the other hand, relying on the overlapping representations of the MSP (see overlap between A and C items in Fig. 5i, dCA1) leads to an advantage for the interleaved condition (t(99)=12.17, p<0.001, *B*_10_>1.0e+6; Fig. 5h), as in human performance on the speeded recognition task.

In sum, similar to generic neural network models employing distributed representations (McClelland et al., 1995), TCM predicts an advantage of interleaved exposure for linking ACs. TCM shares with neural network models the use of distributed representations of item associations (mediated by shared context), rendering it similarly sensitive to interference. Unlike generic neural network models and TCM, REMERGE employs an idealized localist representation that is insensitive to the ordering of input presentations and equally supports interleaved and blocked AC inference. C-HORSE employs both kinds of representations: representations in the MSP are overlapping, as those in generic neural network models and in TCM, whereas the TSP hosts pattern-separated representations similar to those in REMERGE. We propose that both representations exist in the hippocampus, as their combination allows us to account for the pattern of behavior we observed empirically.

## General Discussion

Building direct overlap in representations to encode relationships can promote efficient and powerful memory and generalization. In neural network models that adopt these integrated, distributed representations, information must be presented in interleaved order. Does interleaved exposure similarly facilitate the formation of integrated, distributed representations supporting generalization in humans? We previously put forward a model proposing that, on the timescale of one experimental session, the CA1 subfield of the hippocampus can build distributed representations to integrate related inputs (Schapiro et al., 2017). We thus tested this idea using the hippocampally-dependent associative inference task (Bunsey & Eichenbaum, 1996; Preston et al., 2004), assessing participants’ ability to link indirect associations (AC) after interleaved or blocked exposure to directly associated items (AB and BC). We found that, after interleaved learning, participants exhibited an increased capacity to rapidly recognize item relatedness, to efficiently generalize novel information based on learned associations, and to make inferences according to statistical regularities of item associations. These behaviors reflect properties of distributed representations built from interleaved input in neural network models, consistent with the idea that similar representations underlie human behavior.

Models that encode related items using distributed representations support efficient judgments of item relatedness via directly overlapping representations, whereas retrieval-based models require additional processes to support such judgments (Kumaran & McClelland 2012; McClelland et al., 1986). We postulated that if distributed representations are available for learning on a short timescale, interleaving AB and BC pairs across an experimental session would drive the formation of overlapping representations of A and C. These representations would result in higher sensitivity to the AC association in a setting in which participants make rapid judgments without deliberately searching for AC associations. In a scenario with opportunity for additional retrieval-based processing, we would not expect an impact of presentation order. Indeed, a previous study identified no behavioral difference between interleaving and blocking when participants were asked to deliberately infer associations between indirectly related items (Schlichting et al., 2015). As in this prior study and as predicted, in Exp 1 there was no difference between blocked and interleaved conditions in the standard explicit inference task. In an implicit, speeded task, however, there was significant slowing in the response to interleaved ACs—it was more difficult to indicate that these items had not been studied together. Participants also tended to false alarm to the interleaved ACs, believing that they had studied these items together directly. These patterns did not hold for pairs of items that were unrelated but matched on the distribution of time between presentations (foils), indicating an effect of interleaved *order* rather than a simple effect of temporal proximity in the interleaved condition (see below for a discussion of temporal proximity in interaction with integrative encoding). The results suggest that interleaving supports direct AC recognition, whereas blocked associations may require additional retrieval-based processes (though see discussion on inferring encoding vs. retrieval mechanisms below).

In the context of a recognition task, interleaving arguably *impaired* performance, as it led participants to indicate that they recognized pairs of items that were never studied together. But we expect that interleaving should often benefit behavior. If interleaving builds distributed representations, it should afford automatic generalization of new knowledge amongst related entities. In Exp 2, after learning AB and BC associations, participants learned novel arbitrary attributes of some items, and we assessed their ability to generalize such attributes between interleaved and blocked ACs. We found that generalizing novel associations to indirectly related items was superior in the interleaved condition.

Across the first two experiments, interleaved associations benefitted performance in tasks that required relatively rapid, implicit judgments. Are there situations where interleaving would benefit performance even under more explicit conditions? We hypothesized that a statistical learning paradigm (Saffran et al., 1996) would provide such a situation, where direct associations need to be learned over time from graded co-occurrence frequencies. Distributed representations are especially sensitive to this kind of graded statistical information—indeed, our model indicates a complete failure of localist representations in the hippocampus to support this kind of learning (Schapiro et al. 2017). In Exp 3, we exposed participants to objects with the same pair structure, but presented one at a time in a continuous sequence, with each object temporally adjacent to its pairmate. Consistent with our prediction of a qualitative advantage of interleaved associations in this scenario, performance was superior in the interleaved condition during the explicit inference task, with inference no different than chance in the blocked condition. The results suggest that interleaving is essential for forming representations that permit inference in a scenario that requires integrating statistical information across time.

We do not think our tasks are likely to be process pure — integrated and separated representations likely both contribute to all of our tests. We do think there is evidence, though, for a change in the *relative* contribution of distinct underlying processes. We know that all of our tests are sensitive enough to detect a difference between conditions, as each shows robust effects under the specific conditions predicted by our framework. It is difficult to imagine just one underlying process that could predict a positive effect for speeded recognition and a null effect for explicit inference given discretely encoded pairs while also predicting a positive effect in explicit inference given continuously presented pairs.

Simulations contrasting different computational models of associative inference in the hippocampus provided evidence that at least two forms of representation are likely to contribute. We found that an encoding-based strategy using exclusively distributed representations or a retrieval-based strategy using exclusively localist internal item representations rendered a model unable to account for the full pattern of data: TCM (Howard & Kahana 2002; Howard et al., 2009), employing a distributed code, predicts an advantage of interleaving over blocking but fails to account for the possibility of equivalent performance, as in explicit inference. This advantage of interleaving in TCM persists even when temporal delays are shorter on average between blocked pairs than between interleaved pairs. REMERGE (Kumaran & McClelland 2012), a model that is exclusively localist, accounts for the matched performance, but never predicts the observed interleaving advantage. C-HORSE, employing both kinds of representations via different pathways, can account for both phenomena. These qualitatively different results across models are useful in establishing which classes of models are likely to be able to provide a full account of hippocampal-dependent learning and inference: We suggest that models including both separated and integrated representations are likely needed.

There are many domains where interleaving has proven broadly beneficial for learning (Brunmair & Richter 2019), such as in educational settings (Taylor & Rohrer 2010; Samani & Pan 2021) and in motor skill learning (Goode & Magill 1986). There is extensive work on the effects of interleaved and blocked exposure on learning categories of multidimensional stimuli (see Carvalho & Goldstone, 2015 for a review). Though interleaving stimuli from different categories more often benefits learning and generalization (Birnbaum et al., 2013; Kang & Pashler, 2012; Kornell & Bjork, 2008; Noh, Bjork, & Preston, 2021), this is modulated by several factors, such as the kind of stimuli, the structure of the categories, and task instructions. Many of these findings have been attributed to trial-by-trial attentional effects to different features: Exposure to stimuli from the same category back-to-back (blocking) promotes attention to within-category similarities, whereas exposure to stimuli from different categories back-to-back (interleaving) promotes attention to across-category differences (Carvalho & Goldstone, 2015; Carvalho & Goldstone, 2017). Our study was designed to avoid these kinds of attentional effects: Attention to features in adjacent trials was matched across conditions, as back-to-back trials always, in both conditions, contained completely unrelated pairs. The benefit for interleaving in our data is thus more likely to be related to the benefit observed in neural network models, which do not (typically) have these trial-by-trial attentional biases.

One recent category learning paper found better performance under blocked than interleaved conditions, which the authors argued was supported by ‘factorized’ representations that they suggest may be difficult to explain under an attentional account (Flesch et al., 2018). The authors interpret the results as evidence that neocortex may not be as susceptible to catastrophic interference as neural network models would predict. An alternative interpretation, consistent with the current framework, is that orthogonalized representations in areas DG and CA3 of the hippocampus learn factorized representations and thus can support behavior in scenarios where it is not advantageous to integrate across conditions, as was the case in that study.

In addition to the distributed and localist strategies considered above, another influential proposal for how the hippocampus may carry out associative inference is known as ‘integrative encoding’ (Shohamy & Wagner, 2008; Schlichting & Preston 2015). Integrative encoding posits that studying BC, after having studied AB, triggers reinstatement of the AB memory through pattern completion mechanisms, and an overlapping representation of AB and BC is then encoded (which could, in its strongest form, result in a localist ABC representation). Although this strategy, similar to our account, employs overlapping AC representations formed during encoding to support inference, it relies on the episodic encoding and pattern completion mechanisms of DG and CA3. It is not clear whether integrative encoding predicts an advantage for blocked or interleaved presentation. In theory, temporal proximity between related AB and BC presentations during interleaving could facilitate the formation of an overlapping AB-BC representation. It could also allow integration in two directions, through exposure to AB *after* BC in addition to before BC. However, evidence has been presented for stronger integrative encoding with blocking, where stronger AB memory after repeated AB presentations permits more effective reinstatement of AB during BC learning (Schlichting et al., 2015). It may thus be the case that strength is more important than time lag for successful integrative encoding, and suggests that integrative encoding is unlikely to underlie the findings of interleaved advantage in our tasks. It could be that integrative encoding is more likely to occur when participants are intentionally searching for indirect relationships between direct pairs, which was unlikely to be occurring in our tasks, especially in the statistical learning setting in Exp 3. We also know that the interleaved advantage in TCM and C-HORSE does not arise from shorter time lags in the interleaved condition. Still, we cannot rule out the possibility that humans could show the interleaved advantage through integrative encoding that benefits from shorter time lags in some situations. Future work using computational models that implement integrative encoding could fruitfully explore these possibilities.

Several prior imaging and behavioral studies speak to encoding- and retrieval-based strategies and representations that can underlie associative inference. Some have argued that the hippocampus adopts an encoding-based approach, as in integrative encoding (Shohamy & Wagner 2008; Zeithamova et al., 2012; Schlichting et al., 2014; Schlichting et al., 2015), whereas others have argued that inference is supported by retrieval-based sequential activation (Banino et al., 2016; Koster et al., 2018; Barron et al., 2020; de Araujo Sanchez & Zeithamova 2020; Carpenter & Schacter 2017; Carpenter et al., 2021). We demonstrate that repeated, interleaved exposure facilitates the encoding of integrated representations. This is in line with evidence for encoding-based mechanisms often coming from studies that use interleaved exposure (Shohamy & Wagner 2008; Zeithamova et al., 2012), while evidence for sequential, retrieval-based mechanisms come from studies where related pairs are shown in a blocked manner (Barron et al., 2020; Banino et al., 2016) or with limited exposure (Banino et al., 2016; Koster et al., 2018; Carpenter & Schacter 2017; Carpenter et al., 2021). In both our explicit and implicit assessments, we found faster RTs for direct than indirect pairs, which has sometimes been interpreted as evidence for retrieval-based mechanisms (Shohamy & Wagner 2008; de Araujo Sanchez & Zeithamova 2020). However, models learning distributed representations at encoding also predict stronger associations among directly learned associates, which could lead to slower RTs for indirect pairs in the absence of retrieval-based inference. (We do not think that such a model can explain the full pattern of our data, though, because it would always predict a difference between interleaved and blocked conditions in the explicit inference task). In potential tension with our results, an fMRI study (Schlichting et al., 2015) found that blocked exposure promoted integration of related pairs in the anterior hippocampus. However, encoding- and retrieval-based inference strategies could produce similar results in the slow fMRI BOLD signal: through spreading activation, retrieval-based strategies will activate overlapping sets of items for related inputs even if their underlying representations are not integrated (Kumaran et al., 2012; Chen et al., 2021).

Taken together, we provide behavioral and computational evidence that interleaved exposure facilitates behaviors reflecting the formation of integrated, distributed representations. The rapid timescale of the emergence of these behaviors—across one experimental session—demonstrates great potential for further empirical investigation of this powerful form of representation.

## Appendix

### Model descriptions

#### The Temporal Context Model (TCM)

TCM (Howard & Kahana 2002; Howard et al 2009; Fig. 5a) learns distributed representations of input items by associating them with the context in which they are presented. To simulate our task, we adopted a version of TCM designed to model paired-associate learning (Howard et al., 2009). Across 100 simulations, we present a model with a sequence of object pairs generated following the same protocol as in Exps 1 and 2. Items in each pair are presented sequentially in a randomized order. In TCM, each item *α* is represented by a unique one-hot vector *f*_*α*_ in which the unit representing the item has an activity of 1 and activities for other units are 0. During learning, at each timestep *i*, each input item *f*_*i*_ evokes a contextual input 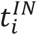 that drives the evolution of a context vector *t*_*i*−1_ according to:

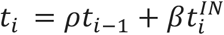

where 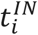 is a combination of a fixed component *c*_*α*_ and a changing component *h* as in Howard et al (2009). If *α* is the item presented at timestep *i*, then 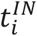 is:

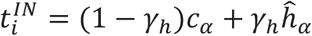

where *c*_*α*_ is identical to *f*_*α*_ and *h*_*α*_ evolves each time *α*is presented according to:

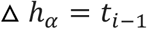

Each time the context drifts, the model updates memory associations between the item and its present context according to:

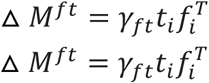

where *M*^*ft*^ and *M*^*tf*^ respectively denote item-to-context and context-to-item associations, and *γ*_*ft*_ and *γ*_*tf*_ are the rates at which these two sets of associations are updated.

Prior to presentation of a new object pair, we present a distractor pattern orthonormal to its current context, which evolves the context at a rate of *β*_*distractor*_. Across learning and at test, to probe the development of item associations, we measured the amount of activation of other items via *M*^*ft*^ and *M*^*tf*^ given an input item. See Supplementary Table 1 for parameter values.

### REMERGE

In simulations of paired associate inference, REMERGE (Kumaran & McClelland 2012; Fig. 5d) represents items (e.g., A and B) and direct pairs (e.g., AB and BC) using a localist code. The model consists of two layers: a feature layer, in which each unit represents a specific item, and a conjunctive layer, in which each unit represents a direct pair. The two layers are connected by bidirectional excitatory connections that link each conjunctive unit with its directly related features units (e.g., the conjunctive unit AB is bidirectionally connected to feature units A and B). The model employs an idealized learning procedure whereby each direct pair presentation strengthens only connections between its conjunctive unit and its constituent feature units (i.e., presentation of pair AB updates only the weights between the conjunctive AB unit and feature units A and B).

At the onset of inference, all units are initialized with an activity of 0. To present a cue item (e.g., A), the activity of its corresponding feature unit is set to 1. Then, activities spread amongst connected units for 150 timesteps starting from *t* = 1.

At each timestep *t* > 1, the net input to a unit *i, net*_*i*_(*t*), is

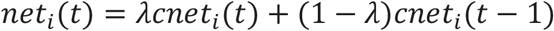

where *cnet*_*i*_(*t*) is the current net input and *cnet*_*i*_(*t* − 1) is the net input at the previous timestep, and *cnet*_*i*_(*t*) is computed as

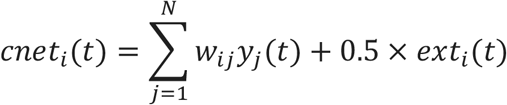

where *y*_*j*_ is the activity of the *j*-th unit connected to the unit *i* and *ext*_*i*_(*t*) is the external input for the unit *i*, which is 1 for the cue item and 0 for all other items across all timesteps.

For a feature unit *i*, its activity *y*_*i*_ is computed as a logistic function of *net*_*i*_, such that:

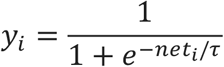

where *τ* is a temperature parameters controlling the degree to which *y*_*i*_ varies with *net*_*i*_.

For the conjunctive unit layer, a hedged softmax function is applied over all conjunctive units to determine the activity of each unit *i*, such that

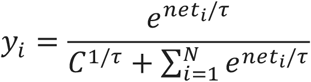

in which *τ* is a temperature parameter, and C is a constant that controls the total activity across conjunctive units. At each timestep, we record activities of all feature and conjunctive units. See Supplementary Table 2 for parameter values.

### C-HORSE

This neural network model of the hippocampus was originally developed in Emergent v.7.0.1 in C++ (Aisa et al., 2008). We ported the model to Golang Emergent v.1.0.5 (https://github.com/emer/emergent), which required the parameter changes detailed in Supplementary Tables 3-5. We also split CA1 into two layers for this simulation, consistent with known anatomical projections, which allowed more control over the impact of the separate representations of the MSP and TSP on the output.

The model has units with activity levels ranging from 0 to 1. A unit’s activity is proportional to the activity of all units connected to it, weighted by the value of the connection weights between them, and modulated by inhibition. The model is set up as an autoencoder, tasked with duplicating the patterns presented to EC_in on EC_out. Weights are adjusted to accomplish this task using a combination of error-driven and Hebbian learning. The error-driven component adjusts connection weights such that activity during two “minus phases” becomes more similar to activity during a “plus phase.” The equations for activity dynamics, inhibition, and learning can be found in O’Reilly et al. (2012).

The learning procedure for this hippocampus model is based on differences in projection strengths between subfields at different phases of the hippocampal theta oscillation (Ketz et al., 2013). At the trough of theta (as measured at the hippocampal fissure), EC has a stronger influence on CA1, whereas at the peak, CA3 has a stronger influence on CA1 (Brankačk et al., 1993). For each learning trial, two items are presented to EC_in (for example A and B, represented by the two corresponding input units taking on a value of 1). There is then one minus phase in which EC_in projects strongly to CA1 and CA3->CA1 is inhibited, which corresponds to theta trough. Next, there is a minus phase where CA3 projects strongly to CA1 and EC_in->CA1 is inhibited, which corresponds to theta peak. There is then a plus phase, in which the target activity (A and B on the output layer) is directly clamped to EC_out and allowed to circulate. Activity throughout the network during each of the minus phases is contrasted with activity during the plus phase, and weights are adjusted so that the patterns of local unit coactivity during each minus phase are shifted more towards those of the plus phase.

