## Supplementary material for "Building integrated representations through interleaved learning"

**Supplementary Table 1** - Parameters for TCM simulations

| *parameter* | *description* | *value* |
|---|---|---|
| $\gamma_{fc}$ | the parameter that controls the learning rate for the feature-to-context matrix | 0.1 |
| $\gamma_h$ | the proportion of $t^{IN}$given by $h$ associated with the input item | 0.9 |
| $\beta_{enc}$ | the rate of context drift during encoding produced by each $t^{IN}$ | 0.7 |
| $\beta_{distractor}$ | the rate of context drift produced by each distractor | 0.99 |
| $\gamma_{cf}$ | the rate of update for the context-to-feature matrix | 0.5 |

**Supplementary Table 2** - Parameters for REMERGE simulations

| *parameter* | *description* | *value* |
|---|---|---|
| $w$ | the weight of bidirectional connections | 1.52 |
| $\tau$ | the temperature parameter for the logistic function and the hedged softmax function | 0.4 |
| C | the constant term for the hedged softmax function | 1 |
| $\lambda$ | the proportion of the net input to a unit given by that on the previous timestep | 0.2 |

**Supplementary Table 3** - Layer parameters for C-HORSE, as modified from Schapiro et al. (2017)

| Layer | Parameter | Schapiro et al. (2017) | Current Model |
|---|---|---|---|
| pCA1 | Layer | Does not exist | Added |
| | # Units | Does not exist | 50 |
| | Inhibition | Does not exist | FFFB (Gi = 3) |
| dCA1 | Layer | Does not exist | Added |
| | # Units | Does not exist | 50 |
| | Inhibition | Does not exist | FFFB (Gi = 2.2) |
| CA1 | Layer | Exists | Removed |
| | # Units | 100 | - |
| | Inhibition | kWTA Avg Inhib 0.25 pct/0.7 pt | - |
| DG | Inhibition | kWTA Avg Inhib 0.01 pct/0.9 pt | FFFB Gi = 24 |
| CA3 | Inhibition | kWTA Avg Inhib 0.06 pct/0.7 pt | FFFB Gi = 4.5 |
| EC_in and EC_out | # Units | 8/15/9 | 18 |
| | Inhibition | kWTA Inhib k = 2/0.5 pt | FFFB Gi = 2.0 |

**Supplementary Table 4** - Projection parameters for C-HORSE, as modified from Schapiro et al. (2017)

| Projection | Parameter | Schapiro et al. (2017) | Current Model |
|---|---|---|---|
| Input -> EC_in | Weight range | 0.25-0.75 | 0.8 |
| EC_in -> DG | Learning rate | 0.2 | 0.4 |
| EC_in -> CA3 | Learning rate | 0.2 | 0.4 |
| CA3 -> CA3 | Learning rate | 0.2 | 0.4 |
| | WtScale.Rel | 1 | 2 |
| CA3 -> pCA1/CA1 | Connectivity | 100% | 0.25 |
| EC_in -> dCA1/CA1 | WtScale.Abs | 3 | 1 |
| | Learning rate | 0.02 | 0.04 |
| dCA1/CA1 -> EC_out[a] | Learning rate | 0.02 | 0.04 |
| pCA1/CA1 -> EC_out[a] | Learning rate | 0.02 | 0.04 |
| EC_out -> dCA1/CA1 | Learning rate | 0.02 | 0.04 |
| EC_out -> pCA1/CA1 | Learning rate | 0.02 | 0.04 |
| EC_out -> EC_in | Weight range | 0.49 - 0.51 | 0.7 |
| | WtScale.Abs | 2 | 1 |
| | WtScale.Rel | 0.5 | 1 |

[a] See Supplementary Table 5 for changes in Absolute Weight Scaling.

**Supplementary Table 5** – Weight scale parameters for C-HORSE, as modified from Schapiro et al. (2017)

| Train | | | | | | | | | |
|---|---|---|---|---|---|---|---|---|---|
| **Schapiro et al. (2017)[a]** | | | | | **Current Model[b]** | | | | |
| **Projection** | **Q1** | **Q2** | **Q3** | **Q4** | **Projection** | **Q1** | **Q2** | **Q3** | **Q4** |
| Ecin -> CA1 | 1 | 0 | 0 | 1 | Ecin -> dCA1 | 1 | 0 | 0 | 2 |
| CA3 -> CA1 | 0 | 1 | 1 | 0 | CA3 -> pCA1 | 0 | 1 | 1 | 1 |
| CA1 -> Ecout | 4 | 4 | 4 | 4 | dCA1 -> Ecout | 1 | 0 | 0 | 1 |
| | | | | | pCA1 -> Ecout | 0 | 1 | 1 | 1 |

| Test | | | | | | | | | |
|---|---|---|---|---|---|---|---|---|---|
| **Schapiro et al. (2017)[a]** | | | | | **Current Model[b]** | | | | |
| **Projection** | **Q1** | **Q2** | **Q3** | **Q4** | **Projection** | **Q1** | **Q2** | **Q3** | **Q4** |
| Ecin -> CA1 | 1 | 0 | 0 | 1 | Ecin -> dCA1 | 1 | 1 | 1 | 1 |
| CA3 -> CA1 | 0 | 1 | 1 | 0 | CA3 -> pCA1 | 1 | 1 | 1 | 1 |
| CA1 -> Ecout | 4 | 4 | 4 | 4 | dCA1 -> Ecout[c] | (4)1 | (4)1 | (4)1 | (4)1 |
| | | | | | pCA1 -> Ecout[c] | 1(4) | 1(4) | 1(4) | 1(4) |

**a** No. of cycles per quarter - Q1: 40, Q2: 20, Q3: 20, Q4: 20.

**b** No. of cycles per quarter - Q1: 25, Q2: 25, Q3: 25, Q4: 25.

**c** Symmetric upregulation of Absolute Weight Scale Values (1 vs 4)

### Supplementary Figures

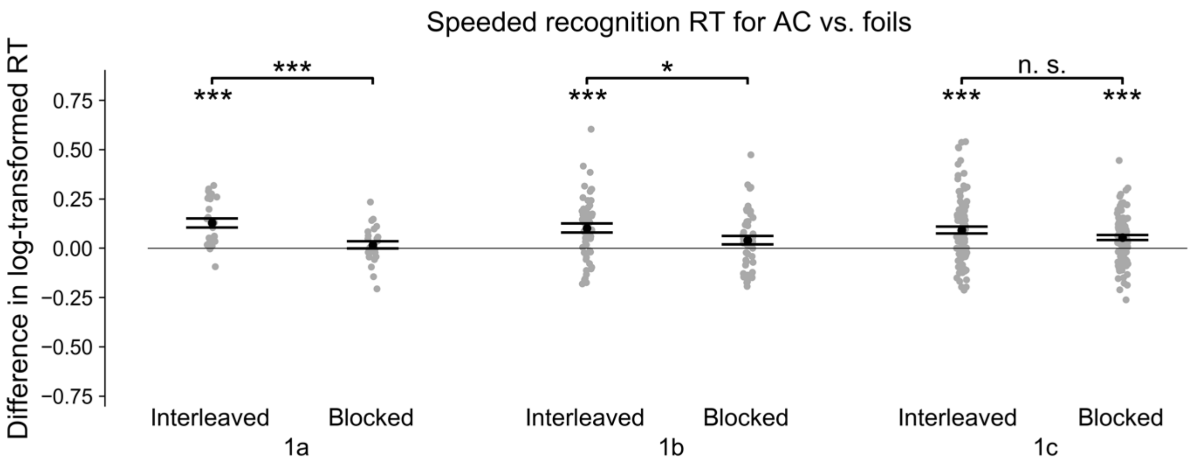


**Figure S1.** RTs for ACs — foils during speeded recognition in Exps 1a-c. *p<0.05; **p<0.01; ***p<0.001.

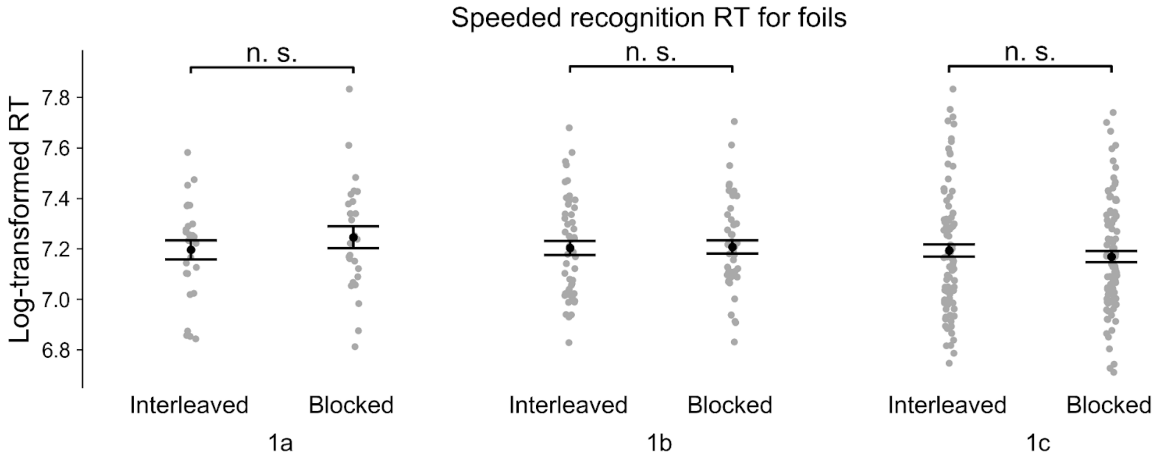


**Figure S2.** RTs for foil trials during speeded recognition in Exps 1a-c.

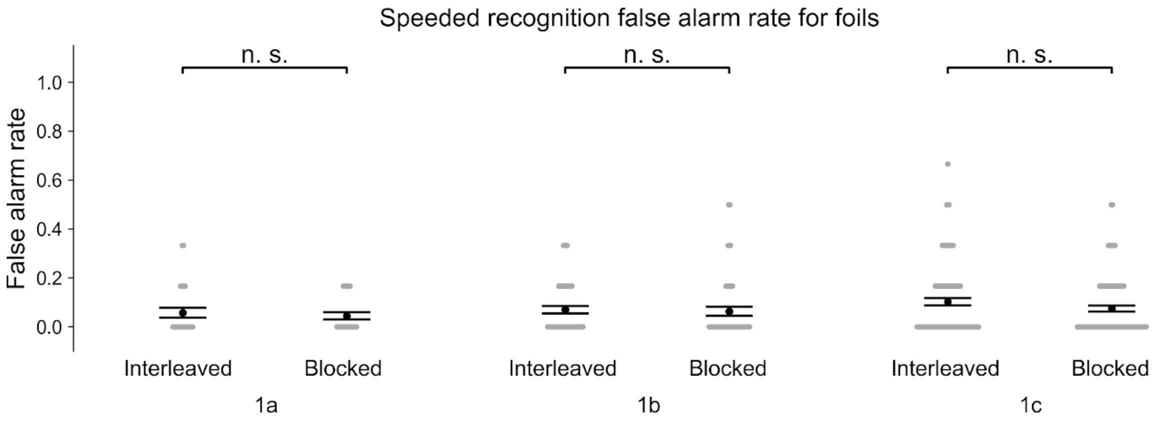


**Figure S3.** False alarm rates for foil trials during speeded recognition in Exps 1a-c.

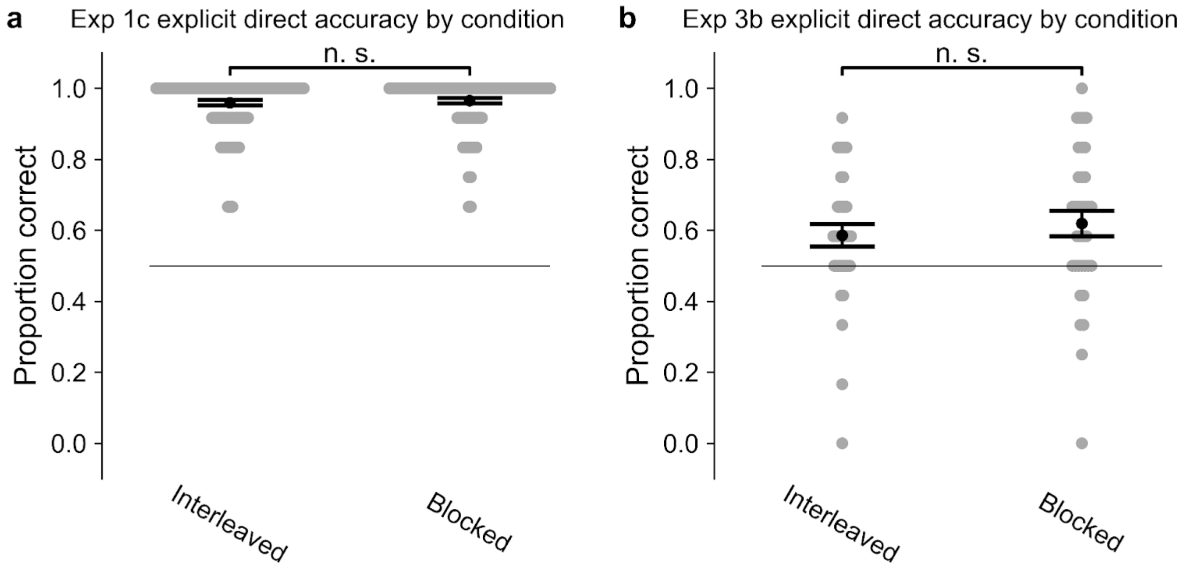


**Figure S4.** Accuracy for explicit direct trials in Exps 1c and 3b.

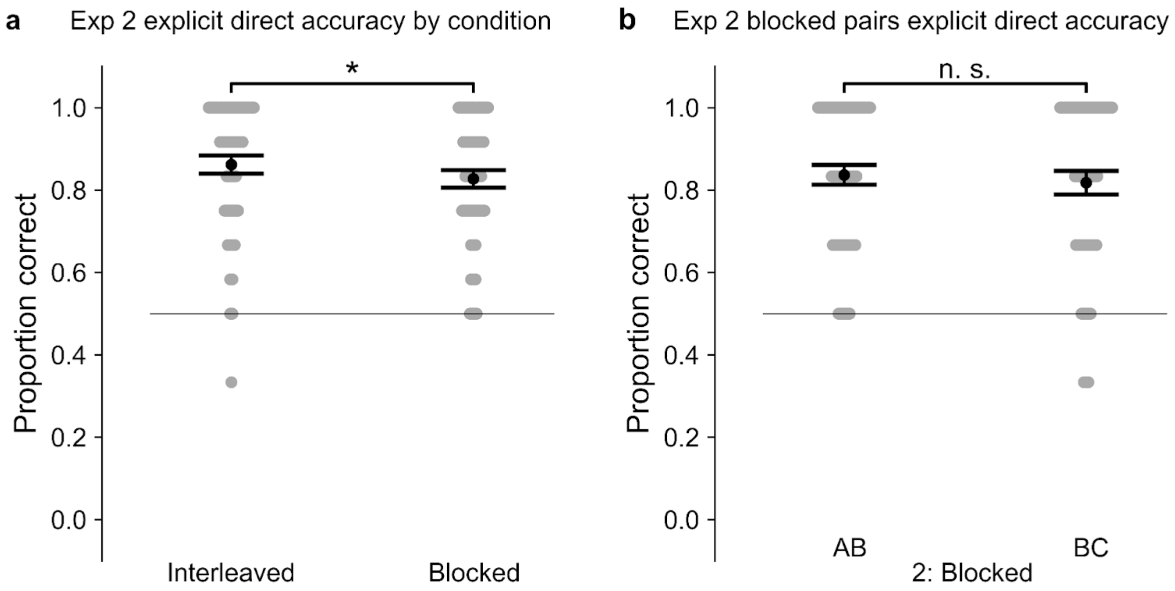


**Figure S5**. Accuracy for explicit direct trials in Exp 2.

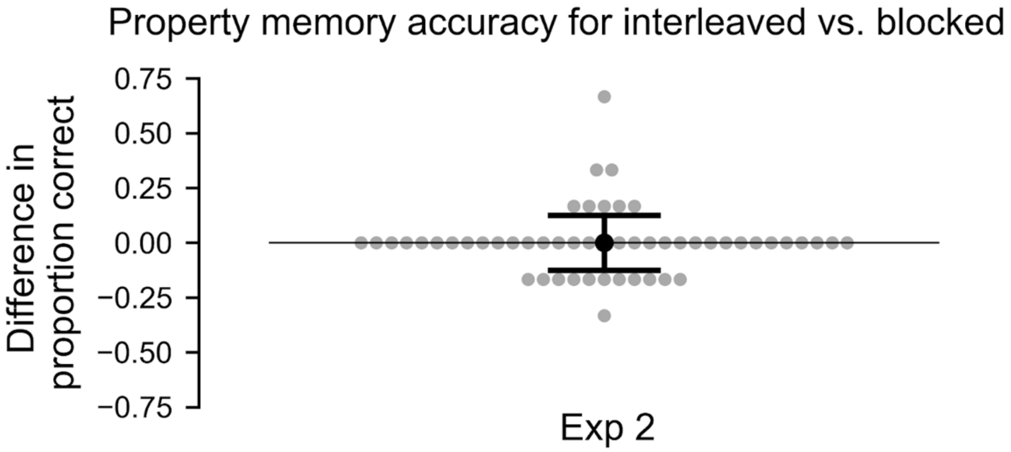


**Figure S6**. Property memory accuracy for interleaved vs. blocked trials in Exp 2.

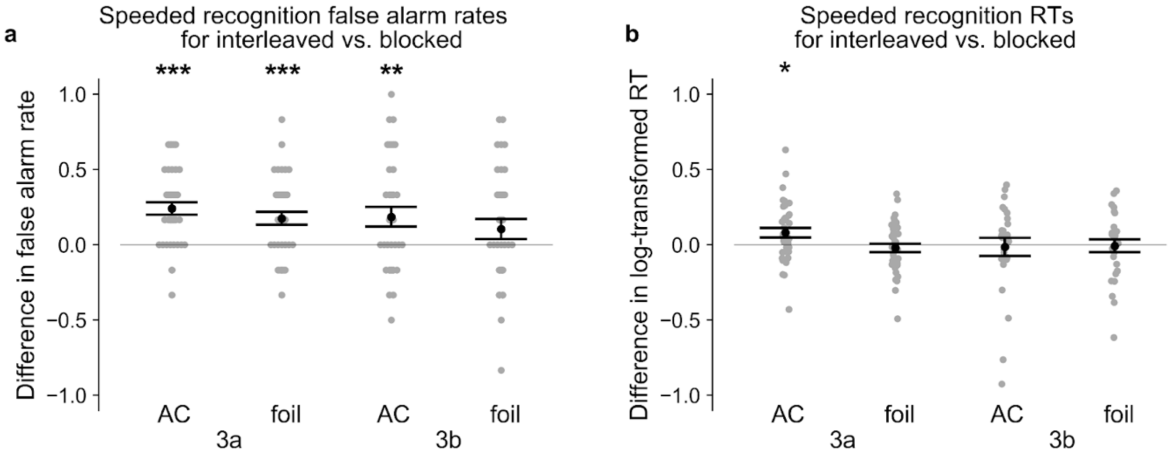


**Figure S7**. (a) False alarm rates for interleaved — blocked AC and foil trials during speeded recognition in Exps 3a-b. (b) RTs for interleaved — blocked AC and foil trials during speeded recognition in Exps 3a-b.

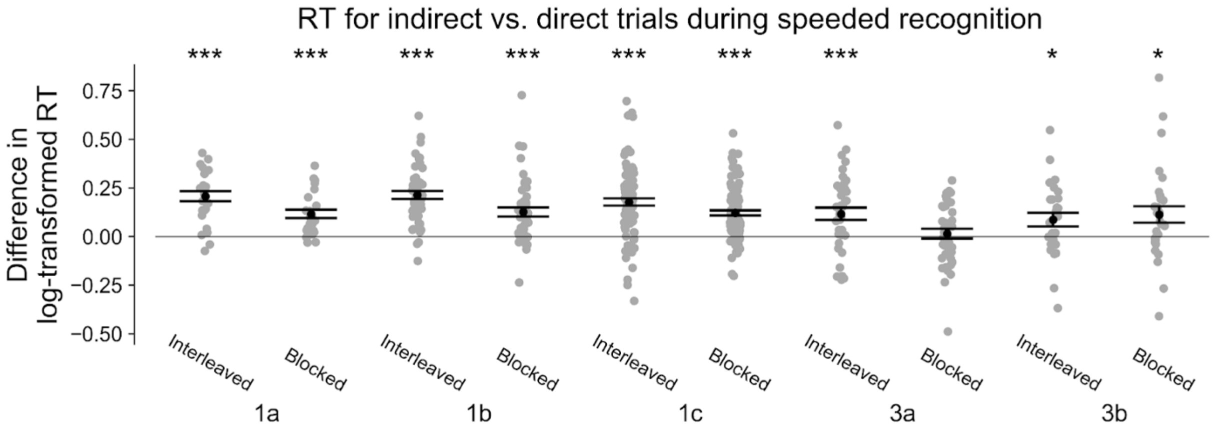


**Figure S8.** RTs for indirect — direct trials during speeded recognition in Exps 1a-c and 3a-b.

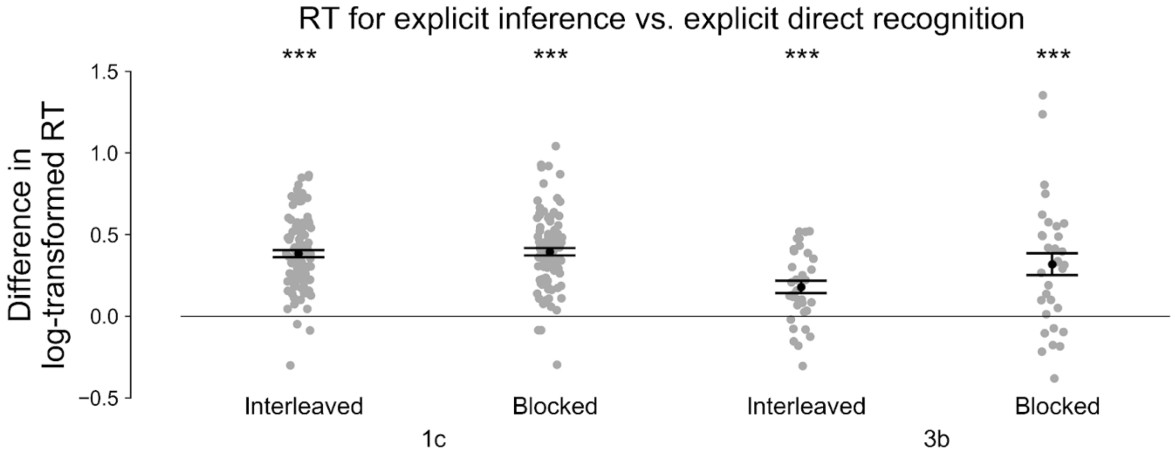


**Figure S9.** RTs for explicit inference — explicit direct recognition trials by condition in Exps 1c and 3b.

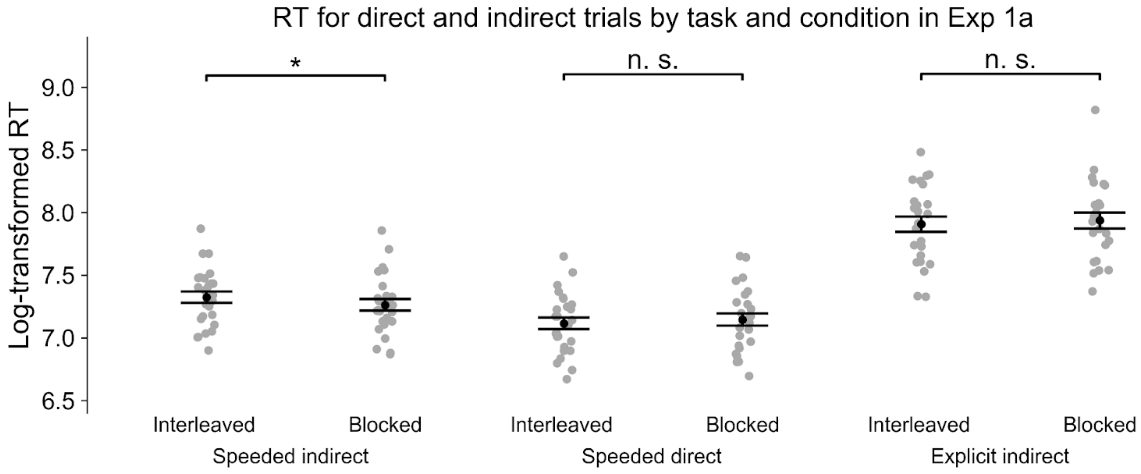


**Figure S10.** RTs for direct and indirect trials by task and condition in Exp 1a.

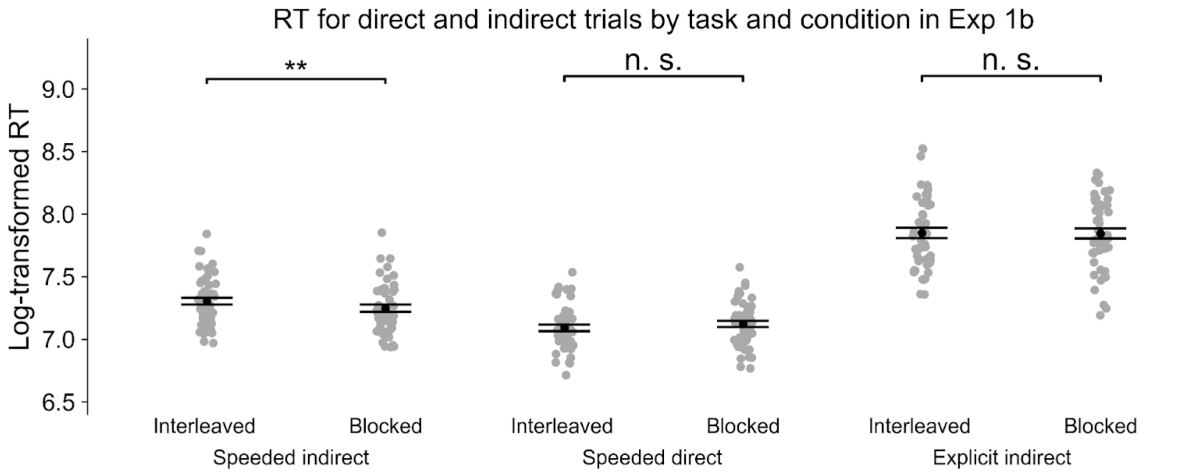


**Figure S11.** RTs for direct and indirect trials by task and condition in Exp 1b.

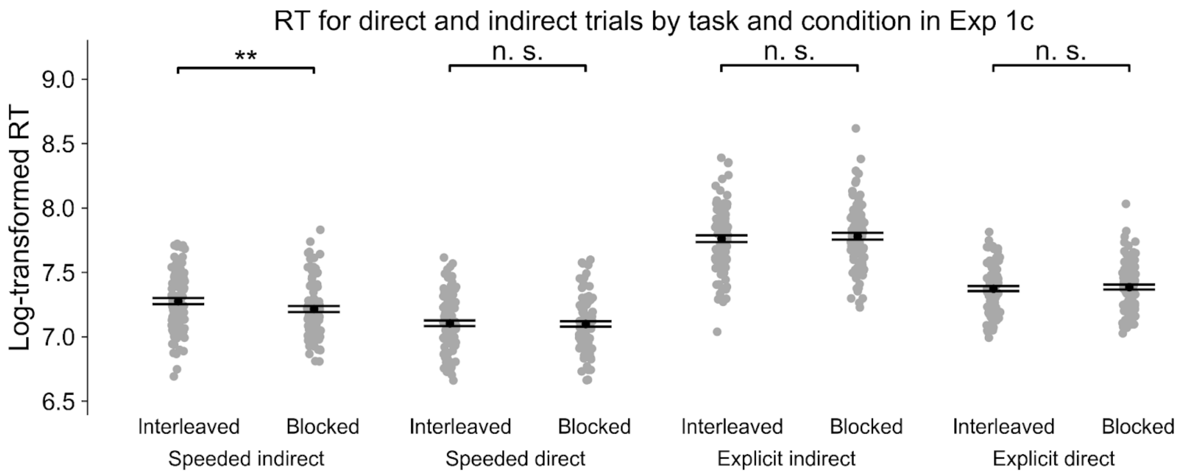


**Figure S12.** RTs for direct and indirect trials by task and condition in Exp 1c.

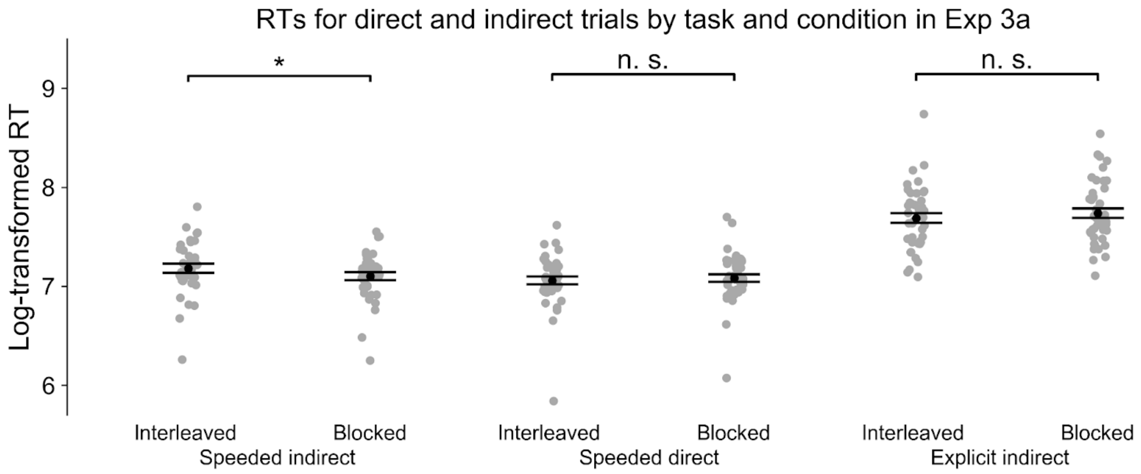


**Figure S13.** RTs for direct and indirect trials by task and condition in Exp 3a.

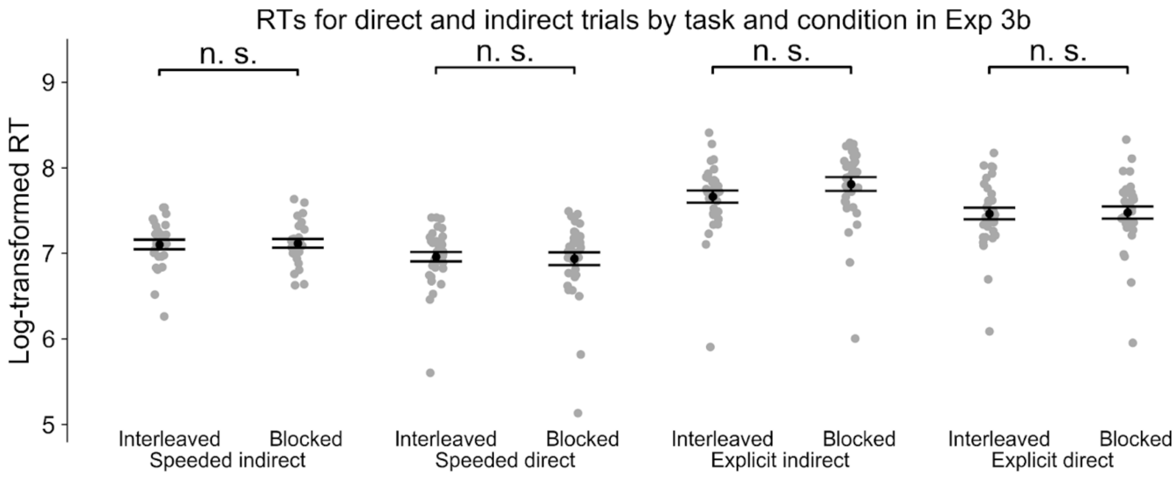


**Figure S14.** RTs for direct and indirect trials by task and condition in Exp 3b.

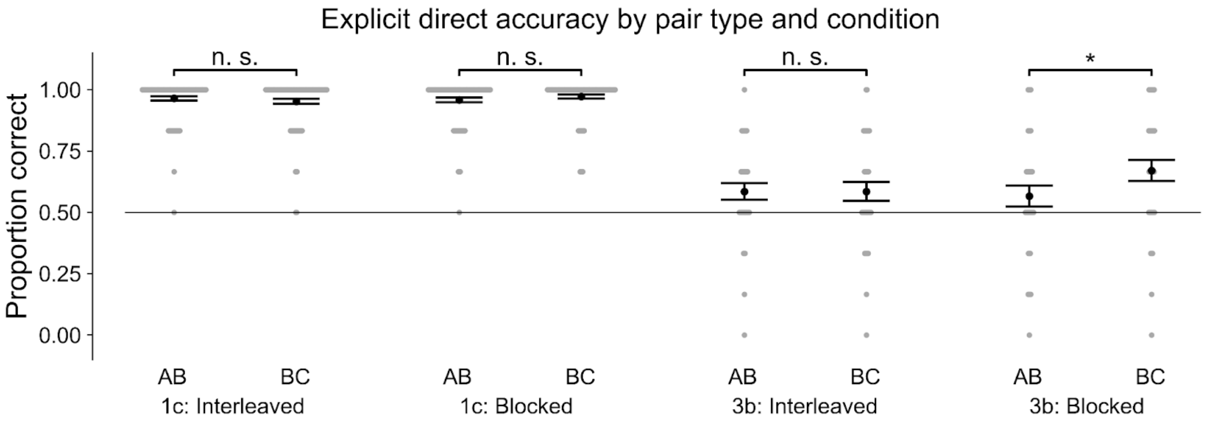


**Figure S15.** Accuracy for explicit direct trials in Exps 1c and 3b.

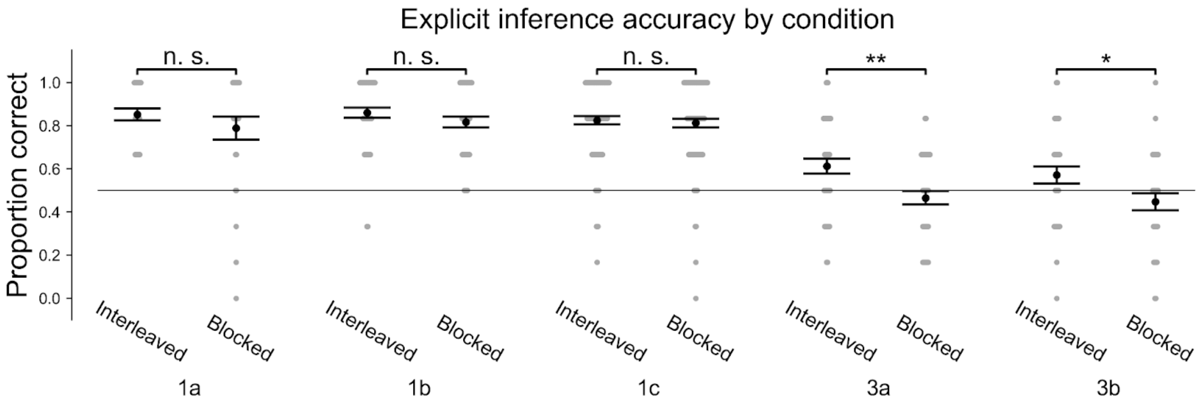


**Figure S16.** Accuracy for explicit inference trials by condition in Exps 1a-c and 3a-b.

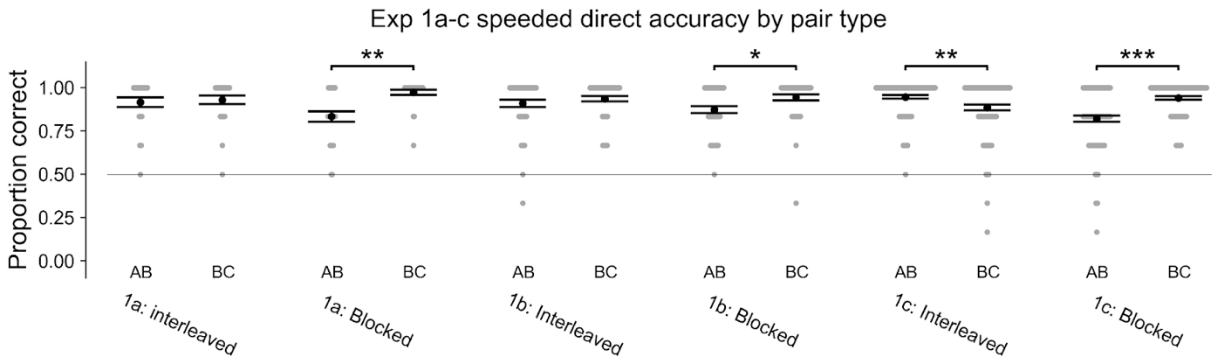


**Figure S17.** Accuracy for speeded direct trials by pair type in Exps 1a-c.

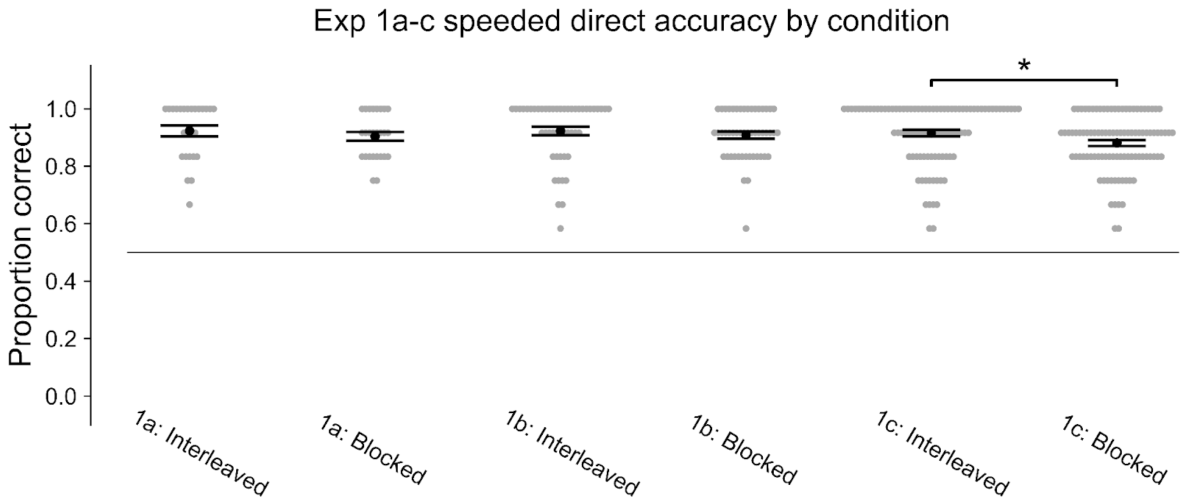


**Figure S18.** Accuracy for speeded direct trials by condition in Exps 1a-c.

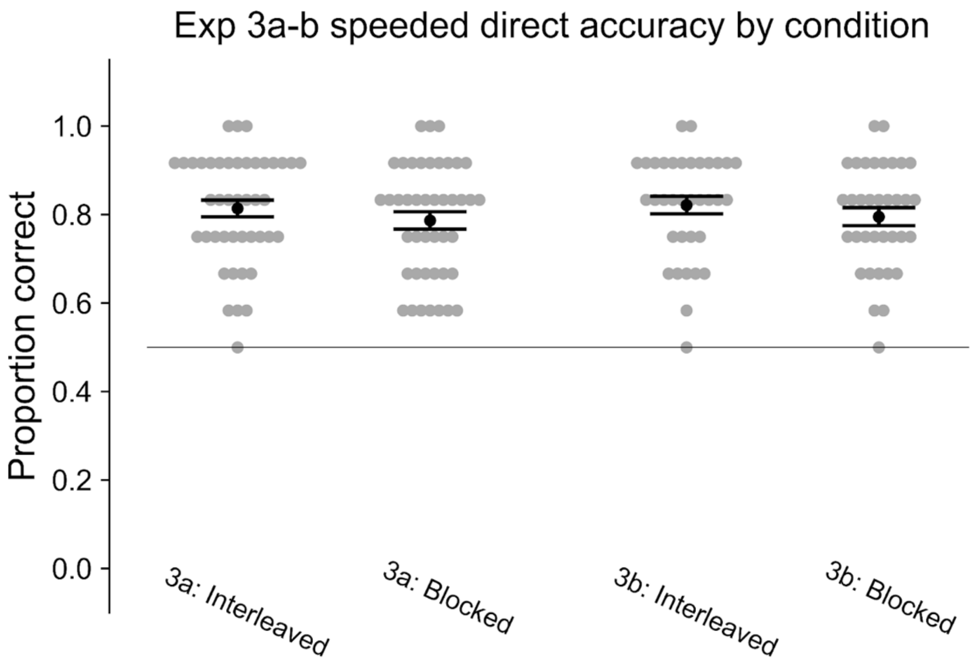


**Figure S19**. Accuracy for speeded direct trials by condition in Exps 3a-b.

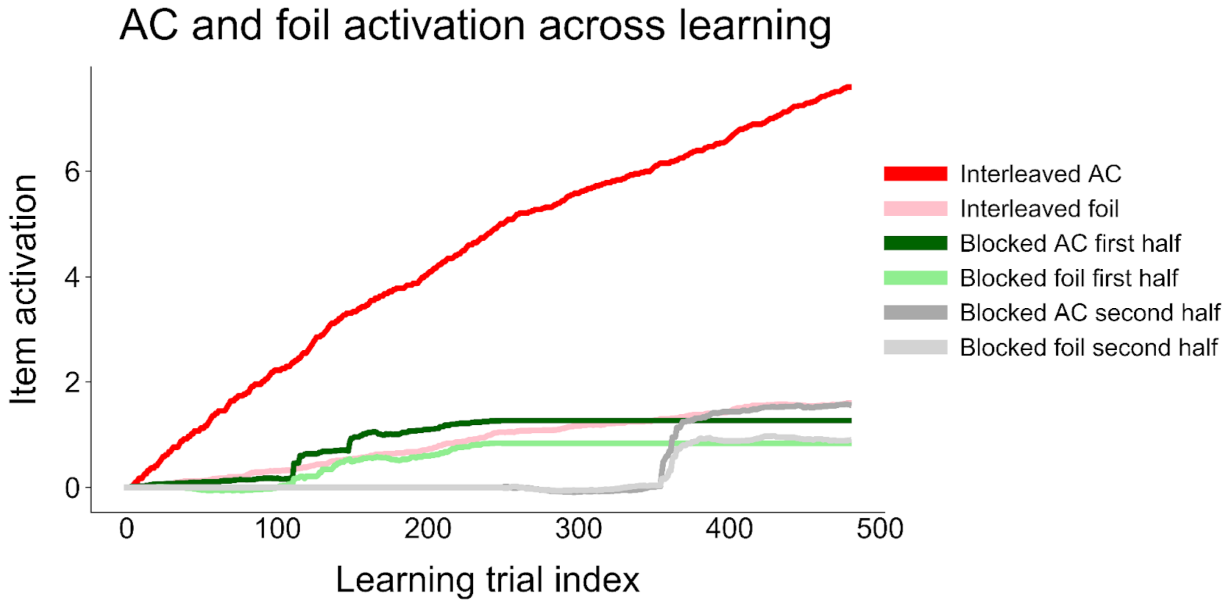


**Figure S20**. Activation of ACs and foils in TCM when related pairs are blocked in the first half of learning, blocked in the second half of learning, or interleaved throughout learning.
